# Unveiling the Epigenomic Control of Temperature Acclimation in Marine Phytoplankton through Multiomics Integration

**DOI:** 10.64898/2026.08.19.745701

**Authors:** Christina Arvanitidou, Marcos Ramos-González, M. Elena García-Gómez, Florence Corellou, Mercedes García-González, Francisco J. Romero-Campero

## Abstract

Temperature plays a central role in marine phytoplankton biogeographical dynamics, physiology and gene expression. Nonetheless, the transcriptional regulatory mechanisms controlling temperature acclimation in marine phytoplankton are yet to be characterized. *Ostreococcus tauri* was chosen as a model species for green marine phytoplankton due to its cellular and genomic simplicity, as well as its evolutionary position within the green lineage. In this study, epigenomic and transcriptomic data were integrated to characterize changes induced by temperature in the trimethylation of histone 3 at lysines 27 and 4 (H3K27me3 and H3K4me3) epigenetic marks established by the Polycomb (PcG) and Trithorax group (TrxG) complexes, respectively. H3K27me3 was found to be a repressive mark responding to temperature, showing predominantly significant increased levels at high temperatures. While H3K4me3 was associated with active transcription, presenting less evident variations in cultures acclimated to different temperatures. H3K27me3 was found only marginally associated with transposable elements, being mostly involved in the repression of specific biological processes, such as gene expression control by transcription factors, meiosis, motors proteins and cytoskeletal structures. No significant conservation was found between the H3K27me3 gene targets in the model plant *Arabidopsis thaliana* and *Ostreococcus tauri*. Nonetheless, transcriptions factors belonging to the MADS-box, WRKY and AP2 families were consistently repressed by H3K27me3 in both species, unveiling that, although the specific downstream targets of this epigenetic mark have diversified during evolution, its role in modulating higher order regulatory nodes remains evolutionary conserved.

## Introduction

Marine phytoplankton are primary producers of essential compounds at the base of marine food webs, and account for approximately 45% of global oxygen production on the planet^1^. Seasonal environmental cues, primarily photoperiod and temperature, play a major role in shaping the dynamic biogeographical distribution of marine phytoplankton^2^. The photoperiodic plasticity of the molecular rhythms at the levels of gene expression, protein abundance and physiology has been extensively characterized using multiomics analysis in marine phytoplankton acclimated to long and short photoperiods^3^. However, despite evidence that temperature significantly influences marine phytoplankton growth^4^, the molecular mechanisms regulating temperature acclimation remains to be explored. This knowledge gap is especially relevant in the current context of climate change, where rising temperatures have been shown to negatively impact marine phytoplankton blooms^5^. Green or chlorophyte marine phytoplankton are evolutionary linked with land plants^6^ in which epigenetic mechanisms, in particular histone post-translational modifications, play a major role in the dynamics of transcriptional regulation in temperature acclimation^7,8^. Among these, the Polycomb group (PcG) marks such as trimethylation of lysine 27 in histone 3 (H3K27me3), and ubiquitination of lysine 121 in histone 2A (H2Aub), as well as the Trithorax group (TrxG) mark, trimethylation of lysine 4 in histone 3 (H3K4me3), have been intensively studied in the model plant *Arabidopsis thaliana*. H3K27me3 is deposited by the Polycomb Repressive Complex 2 (PRC2) and H2Aub by the Polycomb Repressive Complex 1 (PRC1). Both marks have been associated with gene repression, chromatin inaccessibility and local as well as long range genomic interactions^9,10^. Although both marks interact sharing a common set of target genes, H3K27me3 and H2Aub can also act independently marking disjoint set of genes^11^. Temperature has been studied as one of the major environmental cues affecting H3K27me3 dynamics during plant developmental processes such as flowering^12^. In contrast, H3K4me3 is deposited by Trithorax complexes and is associated with gene activation, chromatin accessibility and transcription initiation machinery ^9,13^.

Despite the extensive study of these histone marks in plants and their key role in gene expression regulation, their characterization in microalgae in general and marine phytoplankton in particular, remains in its early stages. Specifically, the genome wide distribution of H3K27me3 has been characterized in the rhodophyte or red microalga *Cyanidioschizon merolae*^14^ and the diatom *Phaeodactylum tricornutum*^15^ where its response to temperature has been reported^16^. In turn, the genome wide distribution of H3K4me3 has been studied in the dinoflagellate *Alexandrium pacificum*^17^ and in the freshwater model chlorophyte or green microalga *Chlamydomonas reinhardtii*^18^ where H3K27me3 remains largely undetectable^19^ with only initial genome-wide studies including other histone marks under nitrogen and sulfur starvation conditions^20^. In addition, comparative evolutionary studies have highlighted an association of H3K27me3 with transposable elements in microalgae^21^.

In this study, *Ostreococcus tauri* has been chosen as a model organism for chlorophyte or green marine phytoplankton due to its genomic^22,23^ and cellular simplicity^24^, ease of cultivation in the laboratory and abundance in coastal marine environments^2^. *Ostreococcus tauri* also occupies an important position in the evolutionary relationships within the green lineage or Viridiplantae^25^. Although the response and acclimation to different light regimes have been extensively studied in *Ostreococcus tauri*^3,26^, research on temperature has only recently been initiated^27–29^. Nevertheless, the molecular mechanisms controlling gene expression during temperature acclimation remain largely unexplored.

In the present work, a phylogenomic analysis identified in the *Ostreococcus tauri* genome the core molecular components of the plant PRC2 and TrxG complexes associated with H3K27me3 and H3K4me3 respectively. In contrast, no component corresponding to PRC1, associated with H2Aub, was found. To investigate the regulatory role of these marks in temperature acclimation in *Ostreococcus tauri* transcriptomic data, RNA-seq (RNA sequencing), and epigenomic data, ChIP-seq (Chromatin ImmunoPrecipitation followed by sequencing) were generated from cultures acclimated to low temperatures, 10°C and 14°C, and high temperatures, 20°C and 26°C. High temperatures led to significant increases in H3K27me3 ChIP levels, associated with gene repression, whereas H3K4me3 showed weaker temperature dependent changes remaining linked to active transcription. H3K27me3 was mainly involved in repressing genes related to cytoskeletal structures, motors proteins, transcriptional regulation and meiosis with only marginal occupation of transposable elements. Although the specific H3K27me3 gene targets were not conserved between *Ostreococcus tauri* and the model plant *Arabidopsis thaliana*, transcription factors from the MADS-box, WRKY and AP2 families were consistently repressed in both species. This indicates that, despise divergence in downstream targets, H3K27me3 role in controlling higher-order regulatory nodes remains evolutionary conserved from green marine phytoplankton to land plants.

## Results

### Identification of PcG and TrxG molecular components in the *Ostreococcus tauri* genome

Four genes encoding histone 3 (H3) were identified in the *Ostreococcus tauri* genome through phylogenomic analysis, Figure 1A. Two of these genes, *ostta05g03770* and *ostta11g02150*, encode the canonical variant H3.1, while the replacement variant H3.3 and the centromeric CenH3 were encoded by single copy genes, *ostta08g00200* and *ostta01g03890*, respectively. A high sequence similarity between these histones and their putative orthologs in other members of the green lineage was observed, Figure 1B. Residues subjected to post-translational modifications such as lysines 4, 9, 27, 36 and 37 were conserved in all proteins except in CenH3. The typical substitutions (A,T)_31_, (S,H)_87_ and (A,L)_90_ and the plant-specific (F,Y)_41_ substitution distinguishing between H3.1 and H3.3 were also identified^30^. The expression profiles in *Ostreococcus tauri* of the genes encoding histone 3 variants is consistent with that observed across highly divergent eukaryotes including mammals and plants^30,31^. Specifically, the two genes corresponding to the H3.1 canonical variant were induced by light and repressed by dark being expressed during the S phase. In contrast, transcripts corresponding to the single gene codifying for H3.3 variant were present during the entire cell cycle with a clear repression by light and activation by dark peaking at the end of the G2 phase, Supplementary Figure 1. This provides strong support for the annotation of *ostta05g03770* and *ostta11g02150* as canonical variant H3.1 and *ostta08g00200* as replacement variant H3.3 in *Ostreococcus tauri*.

**Figure 1.**
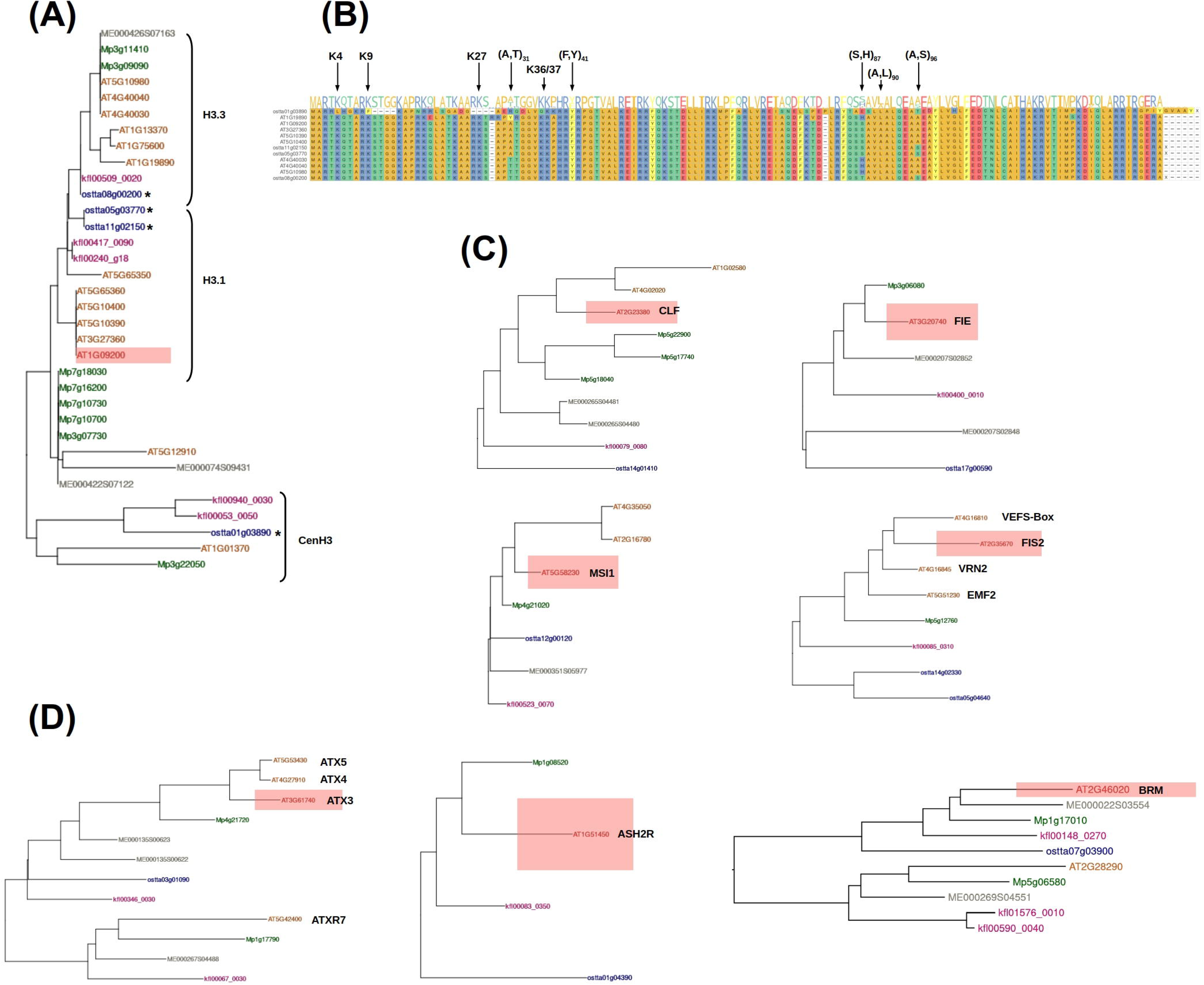
Molecular components of Polycomb Group Repressive Complex 2 (PRC2) and Trithorax Group Complexes (TrxG). **(A)** Phylogenetic tree classifying genes encoding Histone 3 into H3.1 canonical variant, H3.3 replacement variant and CenH3 centromeric variant. *Ostreococcus tauri* genes are represented in blue marked with asterisks, *Arabidopsis thaliana* genes in brown, *Marchantia polymorpha* genes in green and *Mesotaenium endlicherianum* genes in gray. **(B)** Multiple sequence alignment identifying the specific amino acid substitutions characterizing different H3 variants. **(C)** Phylogenetic trees identifying the different components of Polycomb Repressive Complex 2 in *Ostreococcus tauri* as putative *Arabidopsis thaliana* orthologs for CURLY LEAF (CLF), FERTILIZATION-INDEPENDENT ENDOSPERM (FIE) and MULTICOPY SUPRESSOR OF IRA1 (MSI1). **(D)** Phylogenetic trees identifying the different TrxG components in *Ostreococcus tauri* as putative *Arabidopsis thaliana* orthologs for ARABIDOPSIS TRITHORAX 3, 4 and 5 (ATX3, 4, 5); Ash2 RELATIVE (ASH2R) and BRAHMA (BRM). Red rectangles mark the Arabidopsis genes used as representative of the corresponding orthogroup or group of orthologous genes.

Whereas no potential orthologs were found for any of the components of the Polycomb Repressive Complex 1 (PRC1), genes encoding most components of the Polycomb Repressive Complex 2 (PRC2) were identified. Specifically, single copy orthologous genes were found for CURLY LEAF (CLF), FERTILIZATION-INDEPENDENT ENDOSPERM (FIE) and MULTICOPY SUPRESSOR OF IRA1 (MSI1), the *Arabidopsis thaliana* orthologs for Enhancer of zeste (E(z)), Extra Sex Combs (ESC) and Nucleosome remodeling factor 55 (Nurf55), Figure 1C. Two genes were found in the *Ostreococcus tauri* genome encoding orthologs for VEFS genes, the *Arabidopsis thaliana* orthologs for Suppressor of zeste 12 (Su[z]12), Figure 1C. In contrast, no gene encoding for the plant specific PcG proteins LIKE HETEROCHROMATIN PROTEIN1 (LHP1) and EMBRYONIC FLOWER1 (EMF1) were identified. Additionally, no orthologs for the genes involved in PRC2 recruitment in *Arabidopsis thaliana* VIVIPAROUS1/ABI3-LIKE 1 / 2 (VAL1/2)^32^ were found. These genes have also been found to be components of PRC1^33^ further supporting the absence of this complex in *Ostreococcus tauri*.

The components of TrxG complexes have been organized into proteins with H3K4 methyltransferase activity, COMPLEX PROTEINS ASSOCIATED WITH Set1 (COMPASS)-like proteins and ATP-dependent chromatin-remodeling factors^34^. In this respect, single *Ostreococcus tauri* orthologous genes were identified for the methyltransferases ARABIDOPSIS TRITHORAX 3, 4 and 5 (ATX3, 4, 5); the structural components COMPASS ARABIDOPSIS Ash2 RELATIVE (ASH2R) and RETINOBLASTOMA-LIKE PROTEIN (RBL); and chromatin remodelling ATPase BRAHMA and PICKLE, Figure 1D. In contrast, no orthologs for the TrxG co-activators ULT1 and ULT2 were found.

### H3K27me3 and H3K4me3 genome wide distribution at 20°C

ChIP-seq data was generated to characterize H3K27me3 and H3K4me3 genome wide distribution in *Ostreococcus tauri* cultures acclimated to the standard growth temperature of 20°C under constant light.

H3K27me3 occupied approximately 15% of the entire length of the genome, distributed over euchromatic regions in most chromosomes, with no clear accumulation in telomeric or centromeric regions, Figure 2A. Nonetheless, H3K27me3 significantly accumulated in chromosome 19 which has been reported to be associated with viral resistance^35^ and in the first half of chromosome 2 which has been suggested to be involved in sexual reproduction in *Ostreococcus tauri*^36^, Figure 2B. H3K27me3 was mainly located on coding genes with less than 5% of the ChIP signal peaks in intergenic regions, Supplemental Table 1. When associated to genes, H3K27me3 was predominantly found either internal to gene bodies, 43%, or occupying entire gene bodies or only their TSS (Transcription Start Site), 35%, while only marginally associated with TES (Transcription End sites) 17%, Figure 2C. H3K27me3 ChIP levels were significantly different depending on the occupied gene feature, with higher levels when associated to the entire gene body or TSS than when present at the TES or internal to the gene body, Figure 2D. H3K27me3 marked genes exhibited significant low levels of expression compared to non-marked genes, indicating a repressive character of this mark. Genes whose entire body or TSS were H3K27me3 occupied presented stronger transcriptional repression when compared to those in which this mark was internal to the gene body or overlapped with the TES, Figure 2E. H3K27me3 was found marginally associated with transposable elements with approximately only 15% of the marked regions overlapping transposons, Figure 2F. Nonetheless, most H3K27me3 marks located at intergenic regions, 74%, were associated with transposons. Gene Ontology enrichment analysis over the H3K27me3 marked genes revealed that this mark was significantly associated with the repression of specific biological processes, Figure 2G. Namely, H3K27me3 represses gene expression regulation, meiosis and microtubule-based movement, Supplemental Table 2.

**Figure 2.**
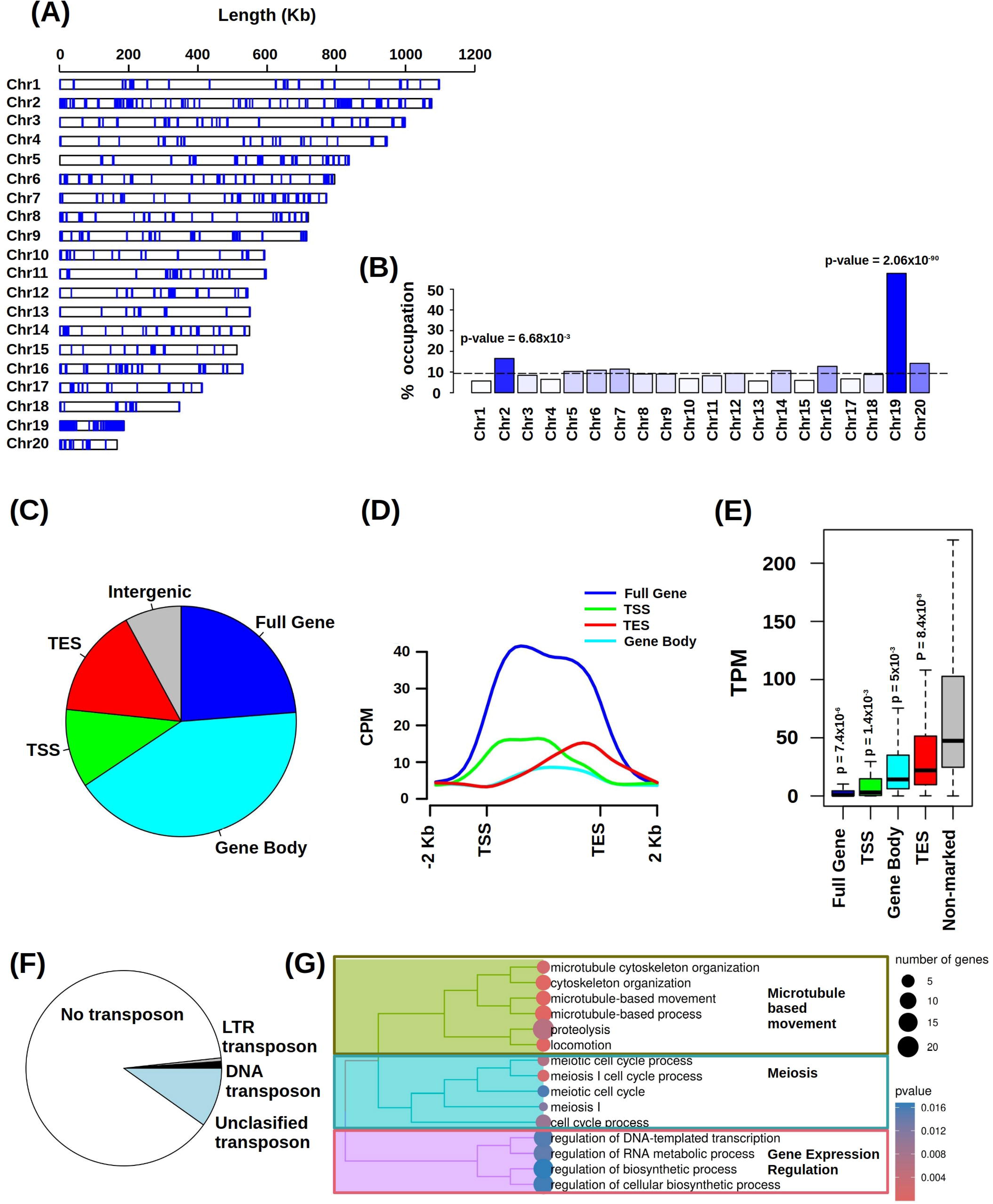
H3K27me3 genome wide distribution and functional annotation in cultures acclimated to 20°C. **(A)** Distribution of H3K27me3 marked regions (blue) over the 20 chromosomes of the Ostreococcus genome. **(B)** H3K27me3 occupation percentage and significance in the different chromosomes. P-values are computed based on normalized Z-scores. **(C)** Fractions of H3K27me3 regions based on the gene feature occupied over marked genes. Full gene body occupation in blue, internal to gene body in light blue, overlapping with the Transcriptional Start Site (TSS) in green, overlapping with the Transcriptional End Site (TES) or intergenic associated to no gene in gray. **(D)** H3K27me3 ChIP signal profile measured using CPM (Counts per Million) over marked genes depending on the location of the mark with respect to gene features. **(E)** Gene expression distribution in the different sets of H3K27me3 marked genes depending on the gene feature occupied. P-values were computed using Mann-Whitney-Wilcoxon nonparametric test. **(F)** Fractions of H3K27me3 regions overlapping with different types of transposons. In white H3K27me3 regions overlapping with no transposon, in gray overlapping with LTR (Long Terminal Repeat) transposon, in black DNA transposons and in light blue unclassified transposons. **(G)** Treemap summarizing Gene Ontology (GO) functional enrichment over the set of H3K27me3 marked genes. GO terms are grouped according to semantic similarities. Node size represents the number of genes and a color gradient from blue to red is used to represent significance.

Specifically, H3K27me3 represses transcription factors involved in gene expression regulation, such as osttaAP2, *ostta14g02090*, and osttaCSP, *ostta02g03030*, Figure 3A. Genes encoding motor proteins associated to long range microtubule based movement, such as minus end directed dyneins osttaDHC1, *ostta06g04460*, and plus end directed kinesins osttaKIN, *ostta18g01040*, were H3K27me3 marked in cultures acclimated to 20°C, Figure 3B. Genes encoding key proteins involved in meiosis were H3K27me3 marked such as Pachytene Checkpoint 2 osttaPA2, *ostta05g02370*, a protein that regulates synaptonemal complex formation ensuring proper chromosome pairing and Disrupted Meiotic cDNA 1 osttaDMC1, *ostta11g01720*, a meiosis-specific recombinase that facilitates interhomologous chromosomal recombination, Figure 3C. Genes involved in lipid metabolism and polyketide biosynthesis, were also H3K27me3 marked under 20°C, such as Enoyl-ACP Reductase osttaENR, *ostta17g00300*, and Polyketide Synthase osttaPKS1, *ostta02g04200*, Figure 3D.

**Figure 3.**
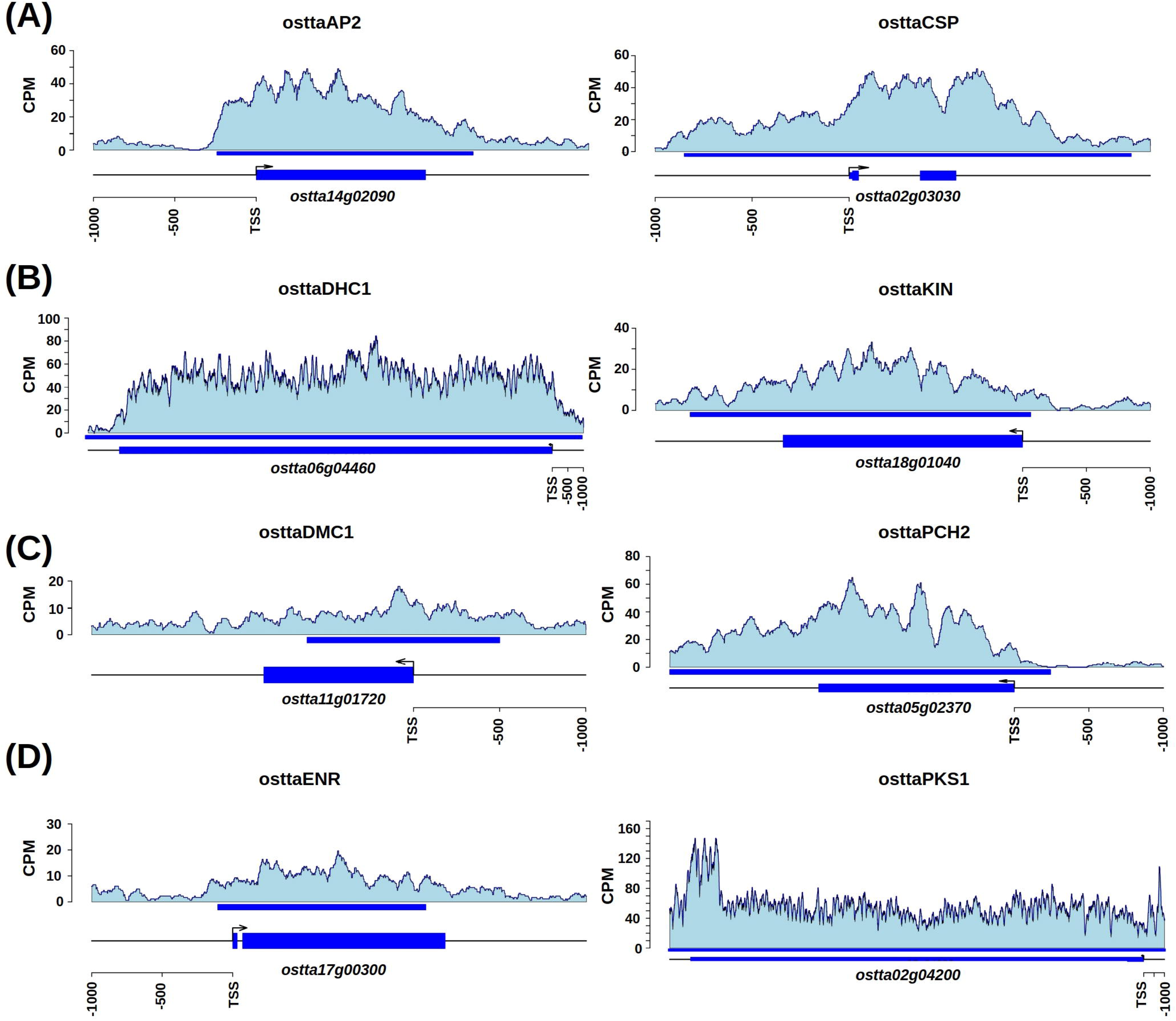
Examples of H3K27me3 marked genes encoding. **(A)** transcription factors belonging to the AP2 (APETALA 2) and CSP (Cold Shock Proteins) families; **(B)** components of motor proteins Dynein Heavy Chain (DHC1) and Kinesin; **(C)** proteins involved in meiosis DNA Meiotic Recombinase 1 (DMC1) and Pachytene Checkpoint protein 2 (PA2); **(D)** enzymes involved in lipid metabolism Enoyl-ACP Reductase (ENR) and Polyketide Synthase 1 (PKS1). H3K27me3 signal profile was measured using CPM (Counts per Million) and represented over marked genes in light blue.

H3K4me3 occupied approximately 10% of the genome length and was distributed across all chromosomes, Figure 4A. Only a slight but significant accumulation of H3K4me3 was observed in chromosome 20, while it was nearly absent from chromosome 19 which was massively marked by H3K27me3, Figure 4B. This distribution indicated that H3K27me3 and H3K4me3 could be mutually exclusive in *Ostreococcus tauri*. H3K4me3 was predominantly associated with genes with less than 5% of its occupancy located in intergenic regions, Supplemental Table 3. This mark was primarily enriched at TSS and along gene bodies, approximately 94% of the marked regions, with less than 2% of the H3K4me3 ChIP signal peaks overlapping only the TES, Figure 4C. H3K4me3 ChIP signal was significantly higher when the marked region occupied the TSS or entire gene body than when located at the TES or internal regions in the gene body, Figure 4D. H3K4me3 was associated with highly expressed genes, presenting significantly higher levels of expression when compared to non-marked genes specifically when located at the TSS or occupying the entire gene body, Figure 4E. In contrast, genes marked at the TES or internal regions of the gene body did not exhibit significantly higher levels of expression when compared to non-marked genes. Functional enrichment analysis of H3K4me3 marked genes revealed significant over-representation of specific biological processes typically associated with high gene expression and active transcription, such as photosynthesis, protein translation, ribosome biogenesis and assembly, Figure 4F and Supplemental Table 4.

**Figure 4.**
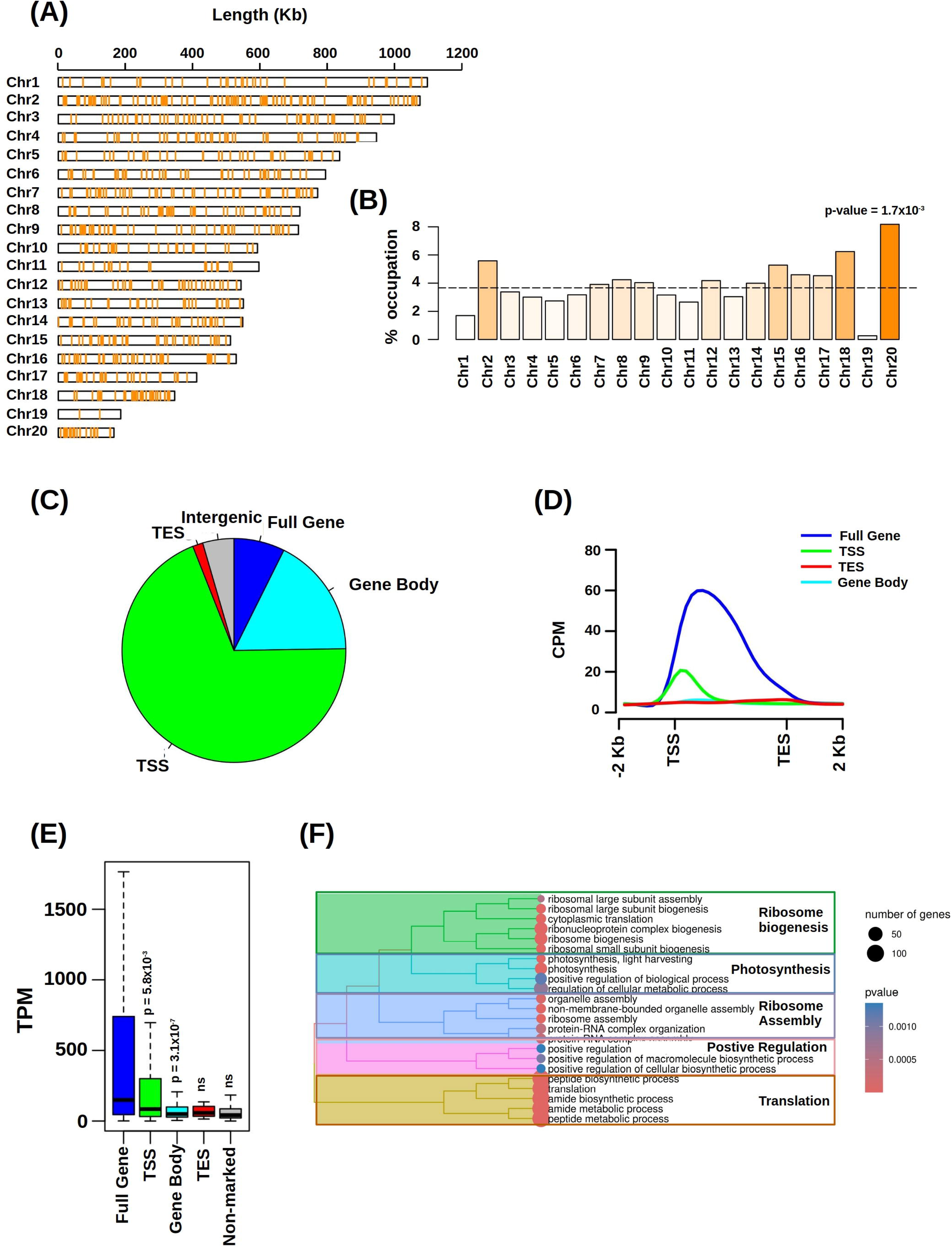
H3K4me3 genome-wide distribution and functional annotation in cultures acclimated to 20°C. **(A)** Distribution of H3K4me3 marked regions (orange) over the 20 chromosomes of the Ostreococcus genome. **(B)** H3K4me3 occupation percentage and significance in the different chromosomes. P-values were computed based on normalized Z-scores. **(C)** Fractions of H3K4me3 regions based of the gene feature occupied over the marked genes. Full gene body occupation in blue, internal to gene body in light blue, overlapping with the Transcriptional Start Site (TSS) in green, overlapping with the Transcriptional End Site (TES) or intergenic associated to no gene in gray. **(D)** H3K4me3 signal profile measured using CPM (Counts per Million) over marked genes depending of the location of the mark with respect to gene features. **(E)** Gene expression distribution in the different sets of H3K4me3 marked genes depending on the gene feature occupied. P-values were computed using Mann-Whitney-Wilcoxon nonparametric test. **(F)** Treemap summarizing Gene Ontology (GO) functional enrichment over the set of H3K4me3 marked genes. GO terms are grouped according to semantic similarities. Node size represents the number of genes and a color gradient from blue to red is used to represent significance.

Specifically, H3K4me3 marked genes involved in ribosome biogenesis and assembly included the large subunit ribosomal protein L3e osttaRPL3, *ostta01g00485*, and the small subunit ribosomal protein S8e osttaRPS8e, *ostta14g02310*, Figure 5A. Genes encoding proteins associated to translation initiation were also H3K4me3 marked, for example, those encoding the translation initiation factors 5A osttaIF5A, *ostta01g03190*, and 3B osttaIF3B, *ostta15g01830*, Figure 5B. Highly expressed genes encoding components of Photosystem II were also H3K4me3 marked, such as those coding for components of the water split complex Photosystem II unit O and P, osttaPsbO, *ostta14g00150*, and osttaPsbP, *ostta14g02630*, Figure 5C. Similar to H3K27me3, H3K4me3 was found marking specific transcription factors such as the single member of the basic helix-loop-helix osttabHLH, *ostta14g01990*, and one of the genes encoding a member of the WRKY family, osttaWRKY, *ostta07g04340*, Figure 5D.

**Figure 5.**
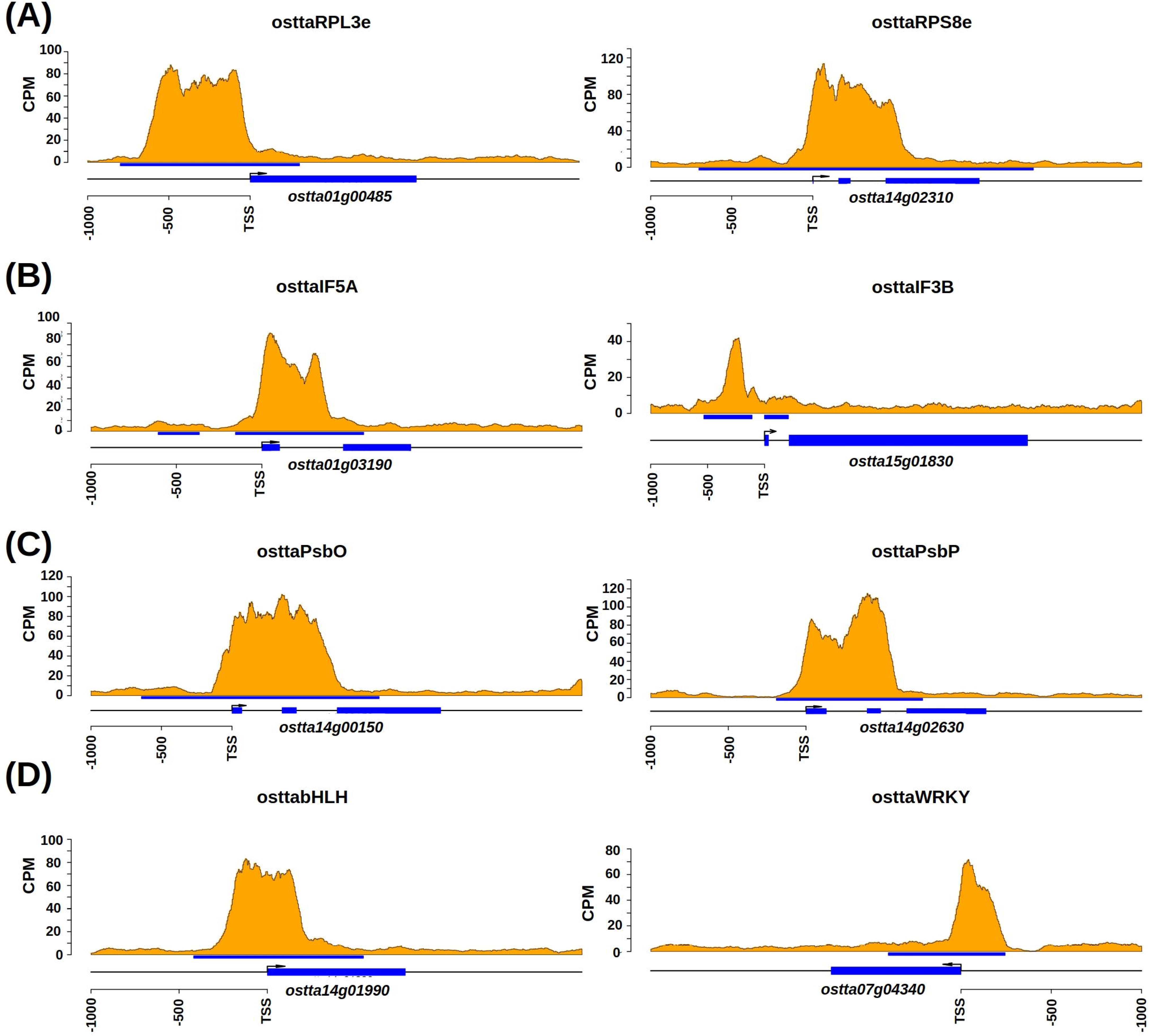
Examples of H3K4me3 marked genes encoding. **(A)** ribosomal proteins RPLe3 (large subunit ribosomal protein L3e) and RPS8e (small subunit ribosomal protein S8e); **(B)** translation initiation factors IF5A and IF3B; **(C)** components of photosystem II PsbO (photosystem II oxygen-evolving enhancer protein 1) and PsbP (photosystem II oxygen-evolving enhancer protein 2); **(D)** transcription factors belonging to the bHLH and WRKY families. H3K4me3 signal profile was measured using CPM (Counts per Million) and represented over marked genes in orange.

H3K27me3 and H3K4me3 were found mostly mutually exclusive, with only 51 genes occupied by both marks, Figure 6A. H3K27me3/H3K4me3 marked genes present significantly lower gene expression than non-marked and H3K4me3 marked genes, indicating a predominant role of the repressive mark H3K27me3 over the mark associated with high gene expression H3K4me3, Figure 6B. The regions occupied by these marks did not overlap suggesting that the same nucleosome cannot carry simultaneously both marks. In the genes with both marks, H3K4me3 was located at the TSS whereas H3K27me3 was found internal in the corresponding gene body reaching in some cases the TES, Figure 6C.

**Figure 6.**
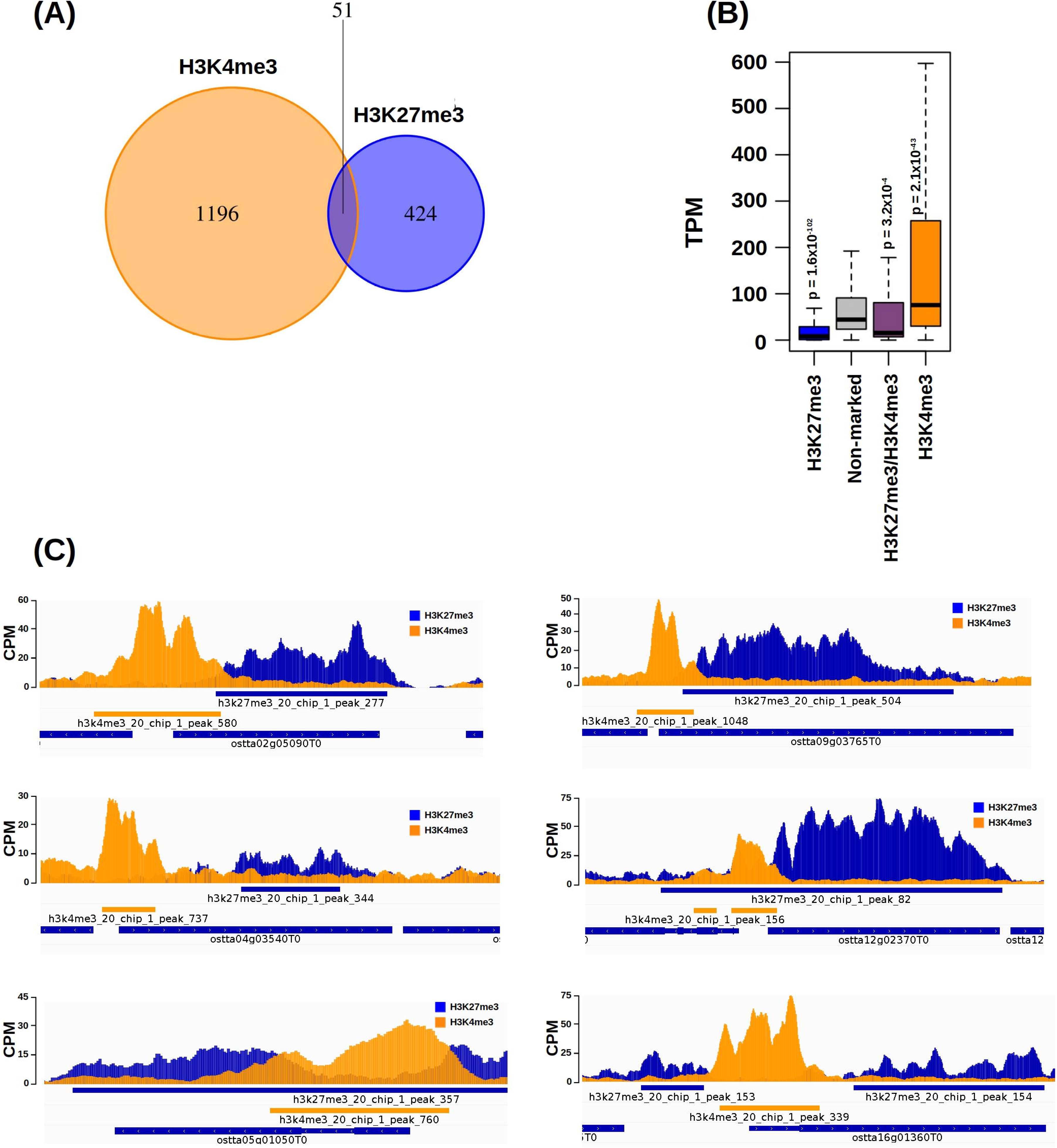
H3K27me3 and H3K4me3 marked genes. **(A)** Comparison of the sets constituted by H3K27me3 marked genes, blue, and H3K4me3 marked genes, orange. **(B)** Gene expression distribution measured as TPM (Transcripts per Million) of the different gene sets, only H3K27me3 marked genes in blue, only H3K4me3 marked genes in orange, H3K27me3 and H3K4me3 marked genes in purple and non-marked genes in gray. P-values were computed using Mann-Whitney-Wilcoxon nonparametric test. **(C)** Examples of H3K27me3/H3K4me3 marked genes. H3K27me3 and H3K4me3 signal profiles were measured using CPM (Counts per Million) and represented over marked genes in blue and orange respectively.

### H3K27me3 significantly changes in temperature acclimation playing a role in gene repression at rising temperatures

H3K27me3 has been reported to play a pivotal role in temperature acclimation and response in plants such as *Arabidopsis thaliana*, regulating developmental processes like flowering^37^. To study the role of H3K27me3 as part of the transcriptional mechanisms underlying temperature acclimation in *Ostreococcus tauri*, the genome-wide distribution of this mark was compared across cultures acclimated to 10°C, 14°C, 20°C and 26°C under constant light in three independent biological replicates.

The genomic regions significantly occupied by H3K27me3 in the three replicates at each temperature were combined to generate a set of temperature consensus H3K27me3 regions. The H3K27me3 ChIP signal was used as an estimation of the occupancy of this mark over the consensus regions in each sample under the different temperatures. Principal components analysis revealed a clear clustering of the different samples according to temperature. H3K27me3 ChIP signals were similar in cultures acclimated to low temperatures, 10°C and 14°C, but clearly different from those acclimated to high temperatures, 20°C and 26°C, which were also different from each other, Figure 7A. The H3K27me3 ChIP signal in the cultures acclimated to 10°C was taken as reference and the number of H3K27me3 regions showing significant changes at 14°C, 20°C and 26°C was determined, Supplemental Table 5. These regions will be referred to as Differentially Occupied Regions (DORs). An increasing number of significant H3K27me3 DORs was observed at rising temperatures, with more DORs showing increased than decreased ChIP signals, Figure 7B. Similar genome-wide accumulations of H3K27me3 at warm temperatures have been observed for the model plant *Arabidopsis thaliana*, being mediated by PRC2 components^38^. These changes were consistent with transcriptomic data in which gene expression was estimated from RNA-seq data generated from the same cultures used to obtain ChIP-seq data. Specifically, principal components analysis revealed a clustering pattern for gene expression similar to that observed for H3K27me3 ChIP signals with samples showing a comparable distribution although a more distinct separation was observed between 10°C and 14°C, Figure 7C. Differentially expressed genes (DEGs) were determined considering 10°C as reference, Figure 7D and Supplemental Table 6. Consistent with the results obtained for H3K27me3, an increasing number of DEGs was detected at rising temperatures with more genes being downregulated than upregulated. This aligns with the repressive nature of H3K27me3 and its predominantly increasing levels at rising temperatures.

**Figure 7.**
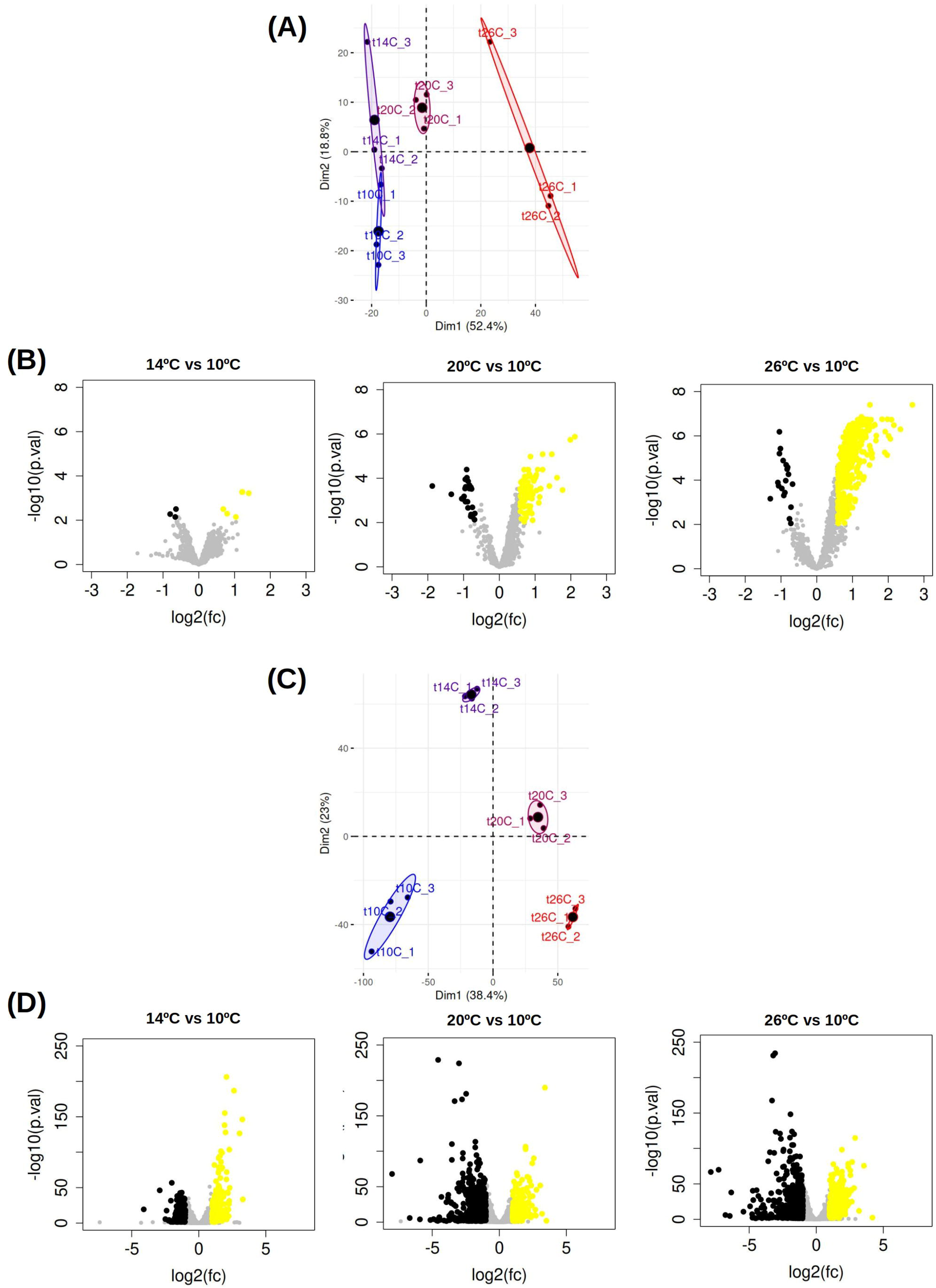
H3K27me3 ChIP signal and gene expression acclimation to temperature. **(A)** H3K27me3 ChIP signal Principal Components Analysis. Individual samples are represented by dots. For each condition a bigger dot is used to represent the average of the three different replicates. Ellipses delimit the 95% maximum likelihood area. Blue represents 10°C, purple 14°C, magenta 20°C and red 26°C. **(B)** Volcano plots representing H3K27me3 Differentially Occupied Regions (DORs) as a response to temperature under 14°C, 20°C and 26°C taking 10°C as reference temperature. H3K27me3 DORs that present reduced levels are marked in black, those that present higher levels are marked in yellow and those H3K27me3 regions that do not change significantly are marked in gray. **(C)** Gene expression Principal Components Analysis in the different samples. Individual samples are represented by dots. For each condition a bigger dot is used to represent the average of the three different replicates. Ellipses delimit the 95% maximum likelihood area. Blue represents 10°C, purple 14°C, magenta 20°C and red 26°C. **(D)** Volcano plots representing differentially expressed genes (DEGs). Repressed or underexpressed DEGs are marked in black, activated or overexpressed DEGs are marked in yellow and genes that do not change their expression significantly are marked in gray.

Gene expression and H3K27me3 ChIP signals were integrated for the genes associated to the H3K27me3 DORs at different temperatures, Figure 8. A significant, gradual and steady increase in H3K27me3 levels was observed at rising temperatures for DORs with increased ChIP signal, Figure 8A. Genes associated with these H3K27me3 DORs presented a significant, gradual and steady decrease in expression levels, Figure 8B, consistent with the repressive nature of H3K27me3. Notably, a significant overlap was detected between the genes downregulated at rising temperatures and those associated with H3K27me3 DORs showing increased ChIP signals, Figure 8C. This suggests that H3K27me3 plays a key role as one of transcriptional regulatory mechanisms underlying gene repression in acclimation to rising temperatures.

**Figure 8.**
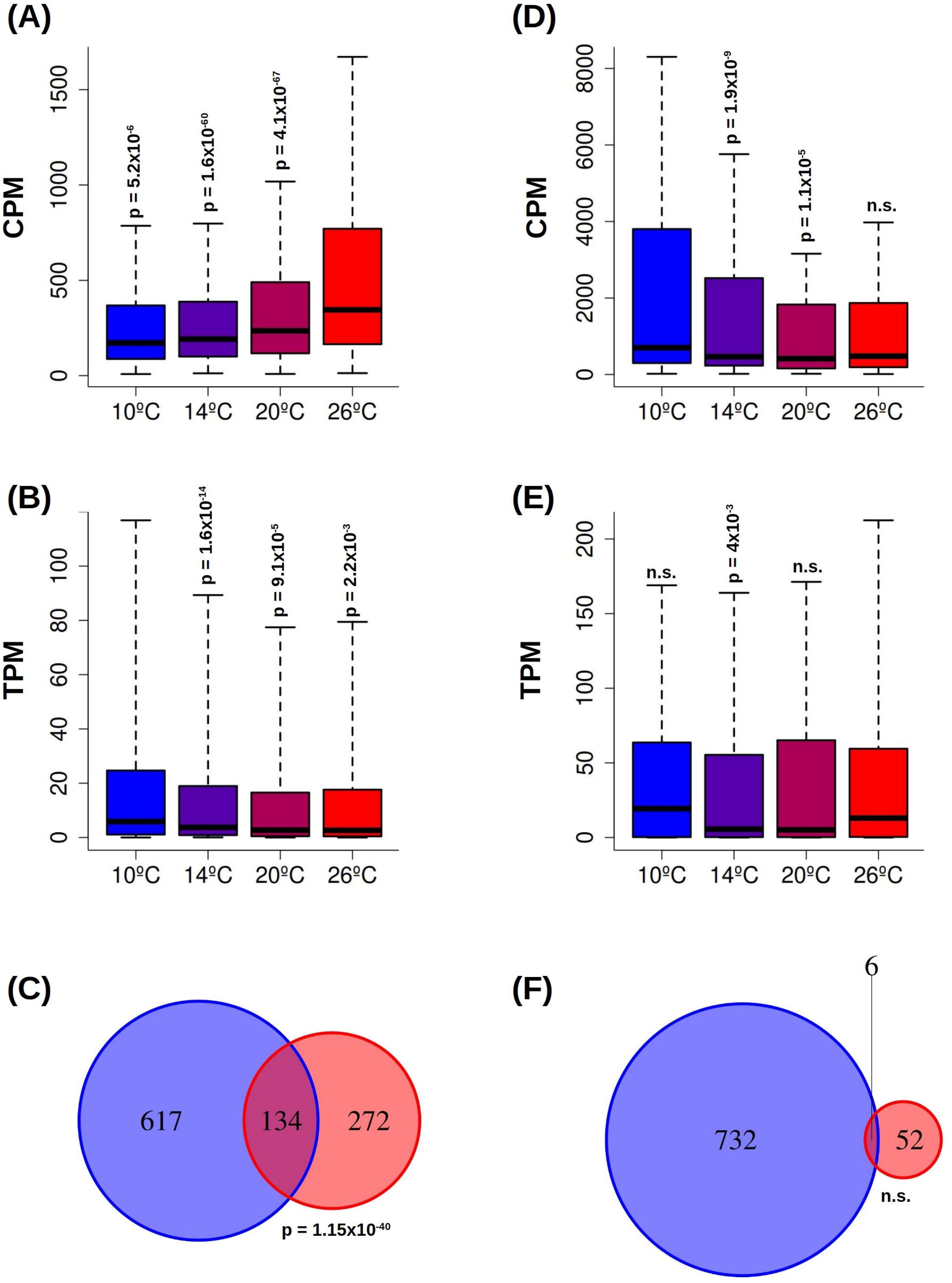
Integration between H3K27me3 and gene expression changes in temperature acclimation. **(A)** H3K27me3 ChIP signal measured as CPM (Counts Per Million) over the marked regions that significantly increase their levels in temperature acclimation. P-values are computed using Mann-Whitney-Wilcoxon nonparametric test. **(B)** Expression level of the genes associated to the H3K27me3 regions that significantly increase their levels in temperature acclimation. P-values are computed using Mann-Whitney-Wilcoxon nonparametric test. **(C)** Venn diagram representing the intersection between the set of differentially repressed genes and the set of genes associated to H3K27me3 marked regions that significantly increase their ChIP signal in temperature acclimation. P-value is computed using a hypergeometric test. **(D)** and **(E)** Similar to A and B for the H3K27me3 marked regions that significantly decrease their levels in temperature acclimation and their associated genes. **(F)** Similar to C for the set of differentially activated genes and the set of genes associated to H3K27me3 marked regions that significantly decrease their ChIP signal in temperature acclimation.

Although a predominant increase in H3K27me3 ChIP signal was detected, some DORs showed decreased levels, Figure 7B. For these regions, a gradual and steady decrease in H3K27me3 levels was observed with rising temperatures, which was significant up to 20°C, Figure 8D. However, this reduction in H3K27me3 ChIP signal did not lead to clear gene activation, Figure 8E. Moreover, no significant overlap was found between the genes upregulated at rising temperatures and those associated with H3K27me3 regions showing decreased ChIP signals, Figure 8E. This is consistent with previous studies showing that the removal of H3K27me3 alone is not sufficient to activate gene expression^9^.

### H3K4me3 associates with highly expressed genes with no clear role in regulating gene expression in temperature acclimation

H3K4me3 has been extensively studied in plants such as *Arabidopsis thaliana* being associated with transcriptionally active chromatin. H3K4me3 has been found strongly associated with the presence of the transcription initiation machinery near the transcription start sites of highly expressed genes. However, different analyses indicate that H3K4me3 does not play a direct regulatory role in gene expression, but rather reflects ongoing transcriptional activity^39^. To investigate the distribution and potential function of H3K4me3 in *Ostreococcus tauri* in temperature acclimation, the genome-wide distribution of this mark was compared across the same cultures acclimated to 10°C, 14°C, 20°C and 26°C used for H3K27me3.

Temperature consensus H3K4me3 regions were defined by combining the genomic regions significantly occupied by this mark in the three replicates at each temperature. H3K4me3 occupancy was estimated using its ChIP signal over the consensus regions in each sample under the different temperatures. Similar to H3K27me3, principal components analysis revealed a clear clustering of the different samples according to temperature with similarities between cultures acclimated to 10°C and 14°C and clear differences between those acclimated to 20°C and 26°C, Figure 9A. H3K4me3 DORs at 14°C, 20°C and 26°C were determined using the ChIP signal in the cultures acclimated to 10°C as reference, Supplemental Table 7. H3K4me3 was found to be less responsive to temperature than H3K27me3 with no significant changes at 14°C and fewer H3K4me3 DORs at 20°C and 26°C than for the case of H3K27me3. Nevertheless, an increasing number of DORs with higher ChIP signal and almost no DORs with lower ChIP signal were observed at rising temperatures, Figure 9B.

**Figure 9.**
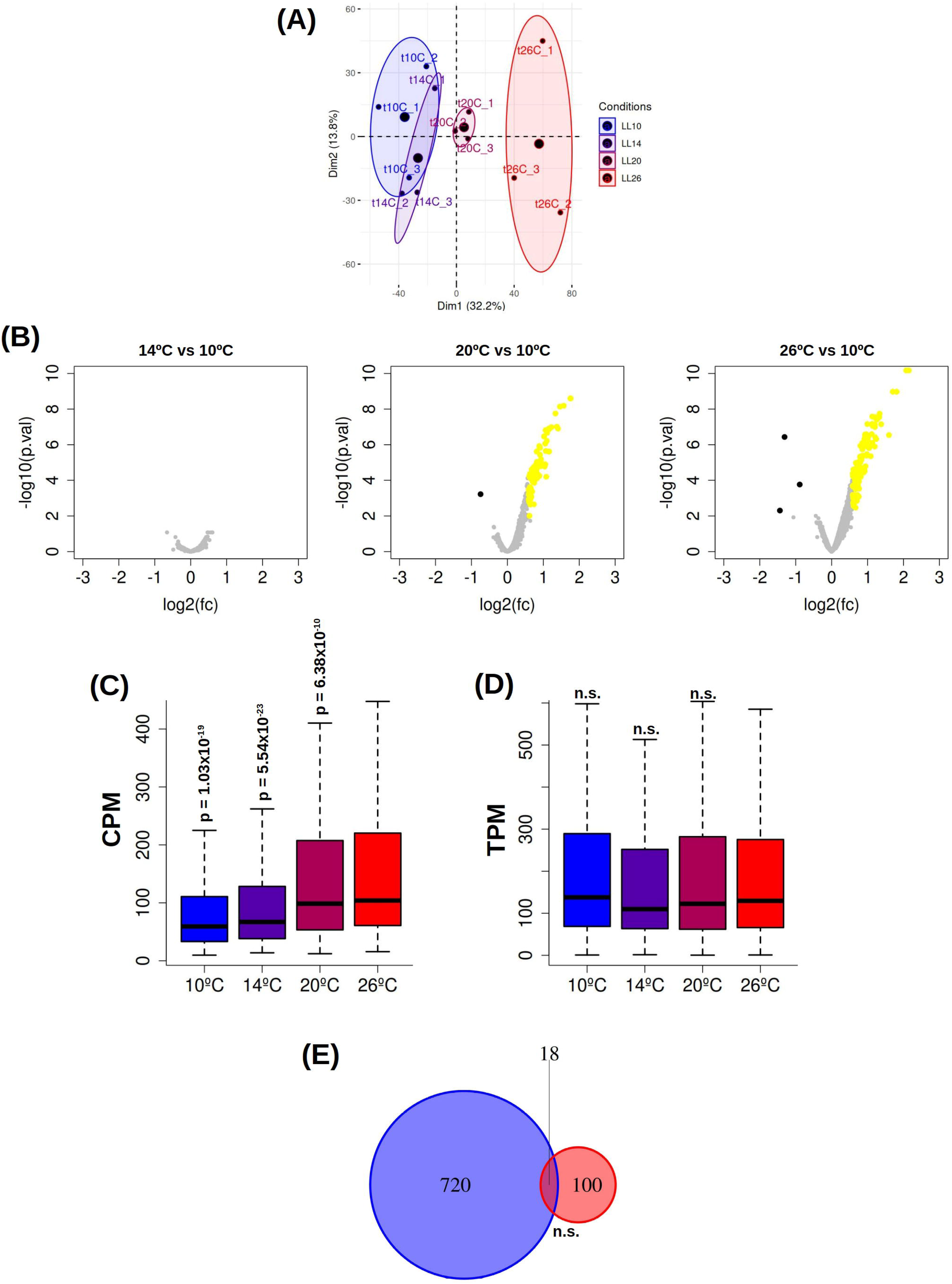
Integration between H3K4me3 and gene expression changes in temperature acclimation. **(A)** H3K4me3 ChIP signal Principal Components Analysis. Individual samples are represented by dots. For each condition a bigger dot is used to represent the average of the three different replicates. Ellipses delimit the 95% maximum likelihood area. Blue represents 10°C, purple 14°C, magenta 20°C and red 26°C. **(B)** Volcano plots representing H3K4me3 Differentially Occupied Regions (DORs) as a response to temperatures under 14°C, 20°C and 26°C taking 10°C as reference. H3K4me3 DORs that present significant reduced levels are marked in black, those that present significant higher levels are marked in yellow and those that do not change significantly are marked in gray. **(C)** H3K4me3 ChIP signal measured as CPM (Counts Per Million) over DORs with increased levels in temperature acclimation. P-values are computed using Mann-Whitney-Wilcoxon nonparametric test. **(D)** Expression level of the genes associated with the H3K4me3 DORs that increase their levels in temperature acclimation. P-values are computed using Mann-Whitney-Wilcoxon nonparametric test. **(E)** Venn diagram representing the intersection between the set of differentially activated genes and the set of genes associated to H3K4me3 DORs with increasing ChIP signal levels in temperature acclimation. P-value is computed using a hypergeometric test.

Although a significant, gradual and steady increase in H3K4me3 levels was observed at rising temperatures for DORs with increased ChIP signal, the corresponding genes did not change their expression significantly with temperature. Moreover, no significant overlap was identified between the genes associated to DORs with increased ChIP signals and activated DEGs at rising temperatures, Figure 9E. This suggests that although H3K4me3 associates with highly expressed genes, its increased ChIP levels do not play a role as a regulatory mechanism underlying gene activation in acclimation to rising temperatures in *Ostreococcus tauri*.

### H3K27me3 regulates cytoskeleton dynamics and lipid metabolism during temperature acclimation

Our data suggest that H3K27me3 plays an important role as a transcriptional regulatory mechanism in temperature acclimation. To elucidate the specific biological processes regulated by this mark, a functional enrichment analysis was performed on the differentially expressed genes (DEGs) associated with H3K27me3 differentially occupied regions (DORs) at rising temperatures, Supplemental Table 8. This analysis revealed the repression of specific biological processes such as cytoskeleton organization, microtubule-based movement and locomotion by motor proteins, Figure 10A.

**Figure 10.**
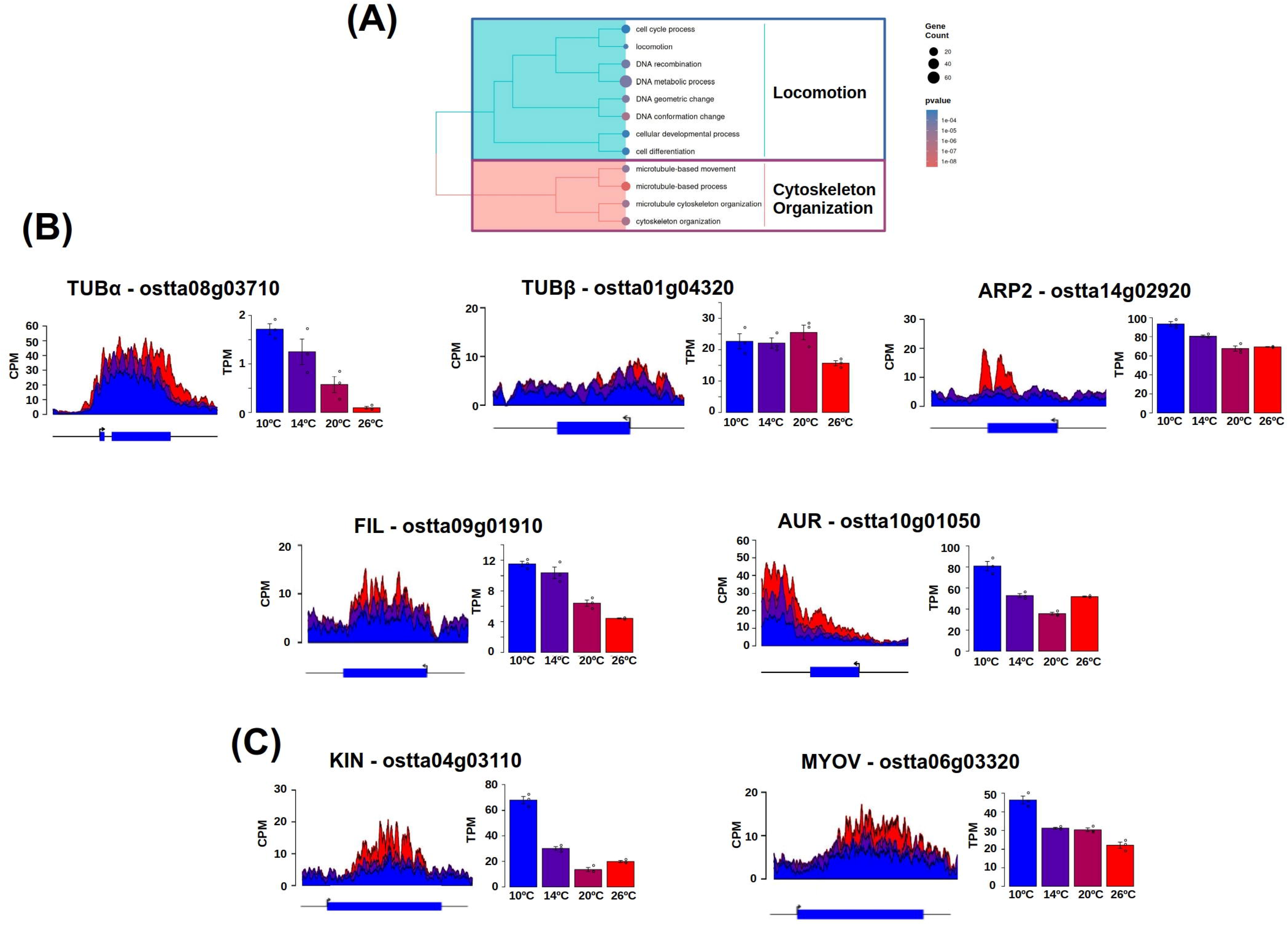
Differentially expressed genes (DEGs) associated with H3K27me3 differentially occupied regions (DORs) at rising temperatures. **(A)** Treemap summarizing Gene Ontology (GO) functional enrichment over the set of DEGs associated with H3K27me3 DORs. GO terms are grouped according to semantic similarities. Node size represents the number of genes and a color gradient from blue to red is used to represent significance. **(B)** H3K27me3 ChIP signal profile measured as CPM (Counts Per Million) and expression barplot measured as TPM (Transcripts Per Million) at rising temperatures from 10°C (blue) to 26°C (red) for genes encoding proteins involved in cytoskeletal dynamics, tubulin alpha (TUBα), tubulin beta (TUBβ), Actin Related Protein 2 (ARP2), MORN-Filamin-Invasin_D3 domain-containing protein (FIL) and Aurora kinase (AUR). **(C)** Similar for genes encoding motor proteins kinesin (KIN) and myosin V (MYOV).

Genes encoding proteins related to cytoskeletal dynamics were expressed at low temperatures and presented increasing H3K27me3 ChIP signals at rising temperatures which was mostly concomitant with gene repression. These include genes involved in microtubule formation such as tubulin alpha (TUBα – *ostta08g03710*) and beta (TUBβ – *ostta01g04320*), cortical actin network establishment such as Actin Related Protein 2 (ARP2 – *ostta14g02920*), cytoskeletal anchoring to membrane like the MORN-Filamin-Invasin_D3 domain-containing protein (FIL – *ostta09g01910*) and kinases involved in microtubule and actin networks stabilization such as Aurora (AUR – *ostta10g01050*), Figure 10B. Concomitantly, at low temperatures high levels of expression were found for genes encoding motor proteins mediating microtubule based transport to outer cellular regions, such as kinesin (KIN – *ostta04g03110*) and actin cortical network based movement for positioning vesicles for secretion such myosin V (MYOV – *ostta06g03320*), Figure 10C.

Transmission Electron Microscopy (TEM) images of *Ostreococcus tauri* cells acclimated to 10°C and 20°C were obtained to investigate whether the temperature-dependent epigenetic regulation of the above mentioned genes could be related to cellular morphological changes. Large vesicle-like structures were observed in cells acclimated to 10°C located in peripheral regions of the cytoplasm, Figure 11A. To further explore the content of these vesicles, cells were stained with the lipophilic dye Nile Red, which specifically binds to neutral lipids. Confocal microscopy images of cells acclimated to 10°C showed, besides the red fluorescence from chloroplasts, distinct Nile Red yellow fluorescence puncta consistent in size and distribution with the vesicles observed in TEM images. This provides evidence of the lipidic content of the vesicles observed at low temperatures, Figure 11B. In contrast, TEM images of cells acclimated to 20°C did not exhibit in large numbers such vesicle like structures, instead they exhibited a large starch granule in the chloroplast. Although some of them presented smaller similar vesicles located in inner regions of the cells, Figure 11C. Consistently, confocal microscopy images of Red Nile stained cells acclimated to 20°C showed mostly only the red fluorescence from chloroplasts, Figure 11D. Accordingly, expression was detected at low temperatures for genes encoding enzymes involved in lipid biosynthesis such as Enoyl-ACP Reductase (ENR – *ostta17g00300*) a component of the fatty acid synthase complex and Diacylglycerol Acyltransferase (DGAT – *ostta16g00910*) involved in triacylglycerols (TAG) synthesis^40^. A similar pattern was detected for the ABC transporter (ABC – *ostta09g01615*) possibly involved in lipid transport as an ortholog of the *Arabidopsis thaliana* gene *ABCG11/WBC11*^41^. H3K27me3 ChIP signal levels increased in all these genes at rising temperatures while gene expression was reduced, Figure 11E.

**Figure 11.**
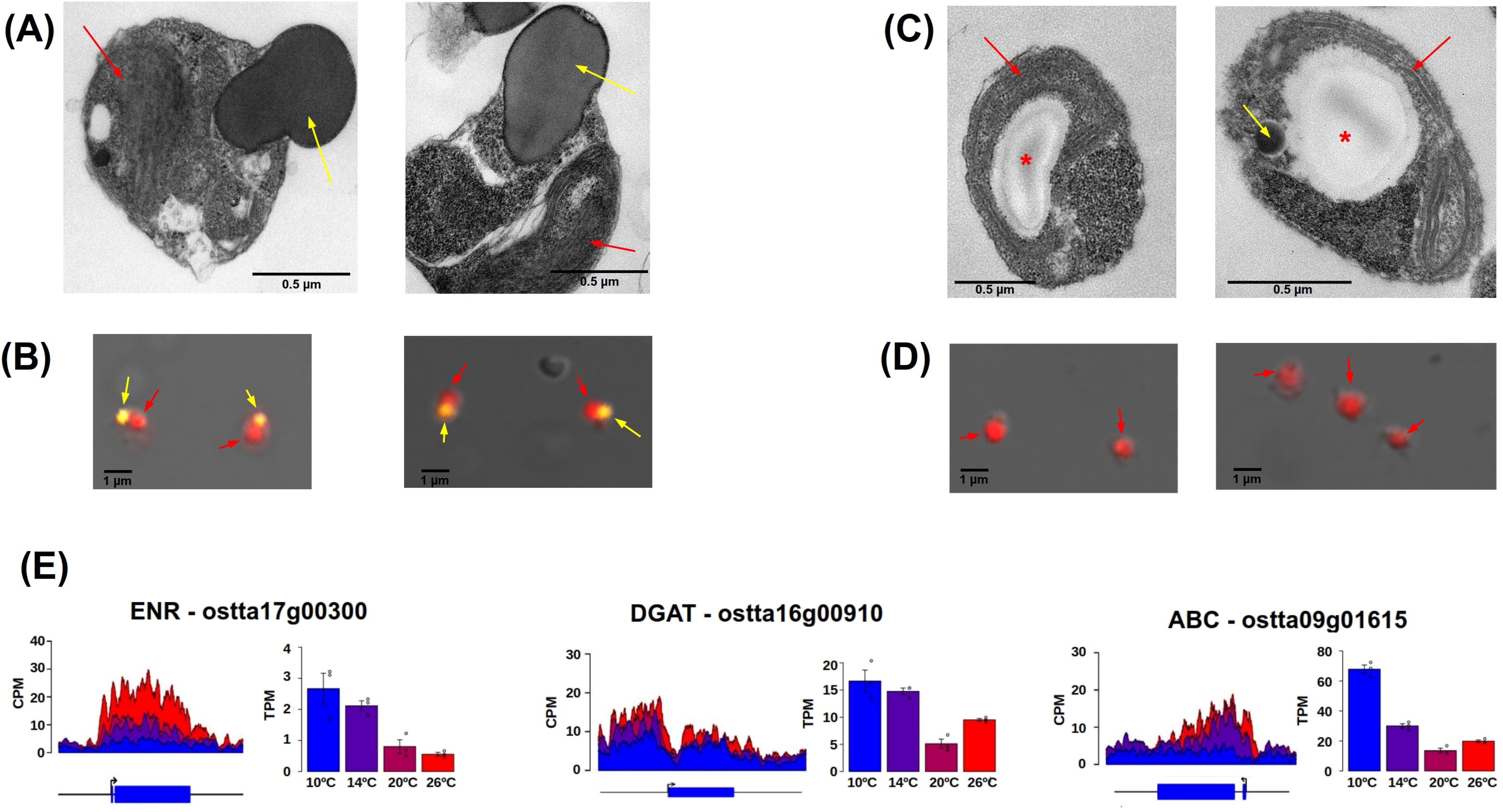
Transmission Electron Microscopy (TEM) and confocal microscopy images of Ostreococcus cells acclimated to 10°C and 20°C. **(A)** TEM images of cells acclimated to 10°C and 20°C. Yellow arrows point to lipid vesicles and red arrows to chloroplasts. **(B)** Confocal microscopy images of cells acclimated to 10°C, Red Nile marked lipid vesicles appear as yellow puncta. Yellow arrows point to lipid vesicles and red arrows point to chloroplasts. **(C)** Similar to (A) for cells acclimated to 20°C. Starch granules are marked with asterisks. **(D)** Similar to (B) for cells acclimated to 20°C. Lipid vesicles are not apparent. **(E)** H3K27me3 ChIP signal profile measured as CPM (Counts Per Million) and expression barplot measured as TPM (Transcripts Per Million) at rising temperatures from 10°C (blue) to 26°C (red) for genes encoding proteins involved in lipid biosynthesis and transport Enoyl-ACP Reductase (ENR), Diacylglycerol Acyltransferase (DGAT) and ABC transporter (ABC).

### H3K27me3 marks the same transcription factor families in *Arabidopsis thaliana* and *Ostreococcus tauri* although no further evolutionary conservation was found

As previously shown, the key molecular components required for the epigenetic mark H3K27me3 identified in *Arabidopsis thaliana*, are also present in *Ostreococcus tauri*. In both species, H3K27me3 predominantly occupies the transcription start site (TSS) and the entire gene body, being associated with transcriptional repression^11^. Moreover, temperature is one of the best characterized environmental cues influencing H3K27me3 dynamics in *Arabidopsis thaliana*^12^ and, in this paper, H3K27me3 has also been shown to play a relevant role in temperature acclimation in *Ostreococcus tauri*. This reveals a conservation of H3K27me3 function between these two evolutionary distant members of the green lineage or Viridiplantae.

To further explore the conservation and divergence in the biological roles of H3K27me3 across these species, the gene sets marked by this post-translational histone modification were compared. Given that the specific genes differ between species and cannot be compared directly, orthogroups or sets of orthologous genes were defined across both complete genomes. An orthogroup was considered to be H3K27me3 marked in *Ostreococcus tauri* when at least one of its member genes from this species was found marked in our data. Similarly, H3K27me3 marked orthogroups in *Arabidopsis thaliana* were defined using previously published data under comparable temperature conditions to those used in this study^11^ .

The H3K27me3 temperature consensus enriched regions identified in this study for *Ostreococcus tauri* corresponded to 994 genes, representing approximately 13% of the entire gene set in its genome. These genes were classified into 749 H3K27me3 marked orthogroups. In *Arabidopsis thaliana*, a greater number of H3K27me3 marked genes were found, 6843 classified into 1859 orthogroups, occupying a larger portion of its genome, 20%. No significant overlap between the H3K27me3 marked orthogroups in each species was found, with only 77 orthogroups being marked in both species, Figure 12A. Nonetheless, functional analysis of the set of the H3K27me3 marked genes in both species revealed a significant enrichment of transcription factors (TFs) involved in gene expression control, Figure 12B. Specifically, members of the AP2, MADS-box and WRKY TF families were found to be H3K27me3 marked in both species. In *Arabidopsis thaliana*, AP2 TF family members such as BABY BOOM (*AT5G17430*), PLETHORA1 (*AT3G20840*) and AINTEGUMENTA (*AT4G37750*), all involved in cell proliferation regulation and stem cell establishment, are H3K27me3 marked^42–44^. In *Ostreococcus tauri*, half of the AP2 TF family members were H3K27me3 marked. Notably, osttaAP2 (*ostta14g02090*) exhibited significant temperature dependent changes in both H3K27me3 ChIP signal and gene expression, suggesting a role in transcriptional regulation in temperature acclimation, Figure 12F. The single MADS-box transcription factor in *Ostreococcus tauri*, osttaMADS (*ostta01g05160*), was also H3K27me3 marked. This gene belongs to the same orthogroup as several well known H3K27me3 marked MADS-box TFs in *Arabidopsis thaliana*, such as FLOWERING LOCUS C (*AT5G10140*), SHORT VEGETATIVE PHASE

**Figure 12.**
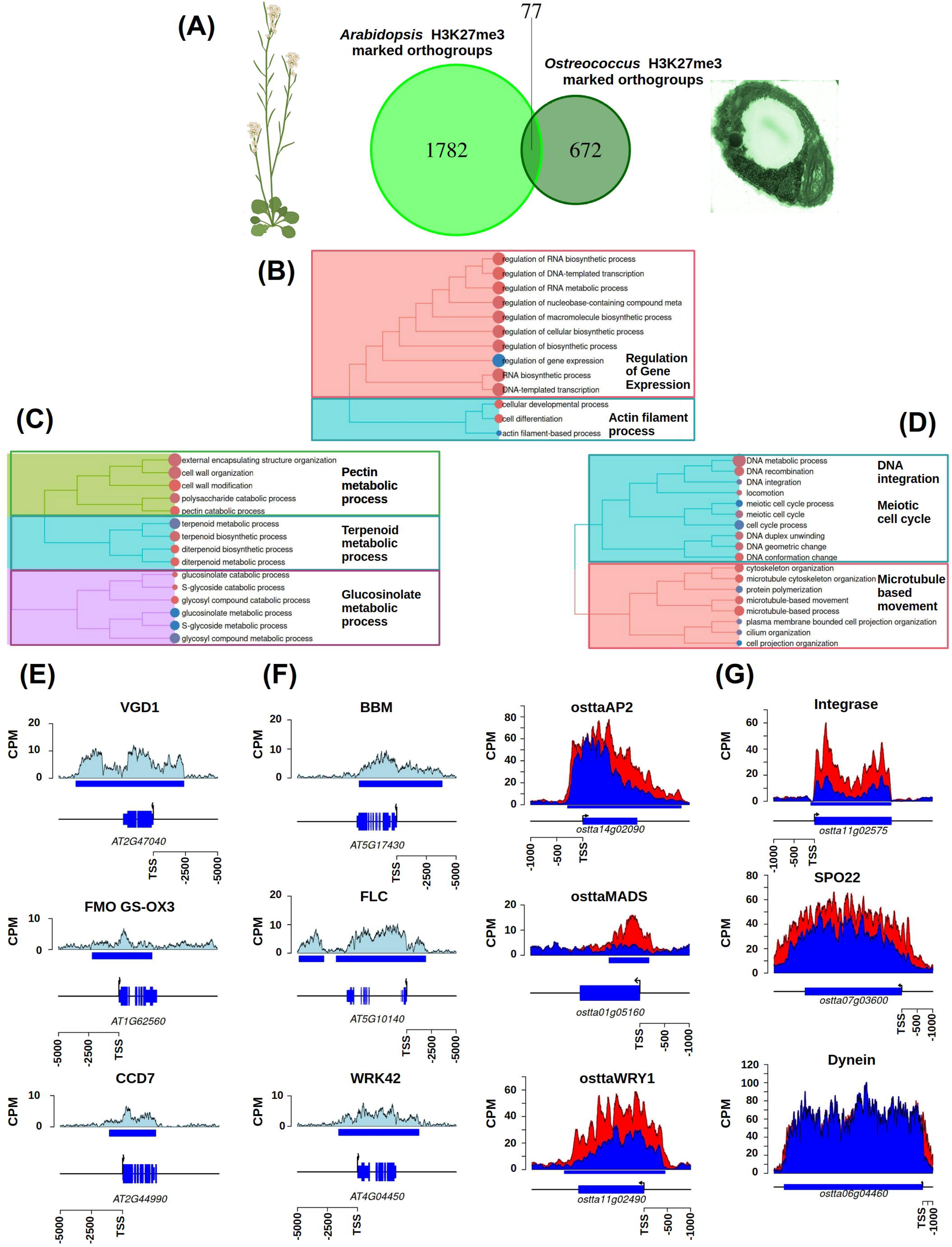
Comparison between the sets of H3K27me3 marked orthogroups in *Arabidopsis thaliana* and *Ostreococcus tauri*. **(A)** Venn diagram representing the common and the species specific H3K27me3 marked orthogroups in Arabidopsis and Ostreococcus. Significance was assessed using a hypergeometric test. **(B)**, **(C)** and **(D)** Treepots representing the functional enrichment analysis over common, Arabidopsis and Ostreococcus specific H3K27me3 marked orthogroups. Semantically similar gene ontology terms are grouped together. The most representative enriched biological processes are identified for each group. **(E)** Examples of Arabidopsis genes associated to the biological processes enriched in the set of Arabidopsis specific H3K27me3 marked orthogroups. **(F)** Examples of Arabidopsis and Ostreococcus orthologuous genes associated to the biological processes enriched in the set of common H3K27me3 marked orthogroups. **(G)** Examples of Ostreococcus genes associated to the biological processes enriched in the set of Ostreococcus specific H3K27me3 marked orthogroups.

(*AT2G22540*) and AGAMOUS (*AT4G18960*), which are involved in floral transition and floral meristem identity^45^, while the role of the *Ostreococcus tauri* MADS-box remains to be characterized. Although H3K27me3 ChIP signal changed significantly in this gene depending on temperature, its expression level was unaffected. Three WRKY TFs were H3K27me3 marked in *Ostreococcus tauri*, osttaWRKY1 (*ostta11g02490*), osttaWRKY2 (*ostta07g04340*) and osttaWRKY3 (*ostta08g02190*). These genes belong to same orthogroup as the Group-II WRKY TFs in *Arabidopsis thaliana* including the H3K27me3 marked genes codifying for WRKY40 (*AT1G80840*) and WRKY18 (*AT4G31800*), which are involved in biotic stress response^46^, and WRKY42 (*AT4G04450*), associated with cold stress response^47^. The specific roles of the corresponding *Ostreococcus tauri* TFs remain to be elucidated, but notably, osttaWRKY1 displayed temperature dependent changes in both H3K27me3 ChIP signal and gene expression, suggesting a possible regulatory role in temperature acclimation.

Orthogroups specifically marked by H3K27me3 in only one of the two species were identified, 672 orthogroups in *Ostreococcus tauri* and 1782 in *Arabidopsis thaliana*, Figure 12A. Approximately half of the *Ostreococcus tauri* specific and one quarter of the *Arabidopsis thaliana* specific H3K27me3 marked orthogroups also contained genes from the other species. This indicates that the species specific marking is not only due to lineage specific gene families, but also reflects differential targeting of conserved gene families. This points to a substantial evolutionary divergence in the genomic targets of H3K27me3 between both species. Indeed, the *Ostreococcus tauri* specific H3K27me3 genes were significantly involved in DNA integration, meiotic cell cycle and microtubule based movement, Figure 12C. The significance of the first process is consistent with the observation that a higher proportion of TEs are H3K27me3 marked in *Ostreococcus tauri*, 13.4%, compared to *Arabidopsis thaliana*, 4%, where H3K9me2/H3K27me1 are the primary histone modifications for TEs^48^. Genes involved in TE activity such as the integrase *ostta1102575,* were exclusively marked by H3K27me3 in *Ostreococcus tauri*, Figure 12G. Regarding meiotic cell cycle, H3K27me3 marked genes exclusively in *Ostreococcus tauri* were found to be particularly associated with the pachytene stage. These include Disrupted Meiotic cDNA 1 (DMC1 – *ostta11g01720*), Pachytene Associated Protein 2 (PA2 – *ostta05g02370*) and meiotic recombination protein (SPO22 – *ostta07g03600*), Figure 12G. The corresponding orthologous genes in *Arabidopsis thaliana* are not marked by H3K27me3, suggesting divergent transcriptional regulation of meiosis between these two species. As described previously, H3K27me3 regulates in a temperature dependent manner microtubule-based movement in *Ostreococcus tauri*. For example, genes encoding for kinesins, *ostta18g01040* and *ostta07g03080,* and dyneins, *ostta14g00070* and *ostta06g04460*, were only H3K27me3 marked in the picoalga. Among these, genes encoding kinesins showed significant changes in H3K27me3 ChIP signals across the temperature range analyzed, whereas genes encoding dyneins remained unchanged, Figure 12G.

Finally, the *Arabidopsis thaliana* specific H3K27me3 genes were significantly enriched in biological processes unique to this species, such as pectin and glucosinolate metabolic pathways, as well as in processes also present in the picoalga, like terpenoids biosynthesis, further supporting the divergence in H3K27me3 targets in these two species, Figure 12B. Pectin is a major component of cell wall in *Arabidopsis thaliana* and it is absent in *Ostreococcus tauri*, which instead possesses a glycoprotein based layer surrounding the plasma membrane. For instance, the pectin methylestarase VANGUARD1 (VGD1 – *AT2G47040*) is H3K27me3 marked in *Arabidopsis thaliana* and lacks an ortholog in *Ostreococcus tauri*, Figure 12E. Similarly, glucosinolates, which play roles in defense and sulfur metabolism, are specific to Brassicaceae species such as *Arabidopsis thaliana*. The flavin-monooxygenase (FMO GS-OX3 – *AT1G62560*) involved in aliphatic glucosinolate biosynthesis is another example of an H3K27me3 marked gene in *Arabidopsis thaliana* without an ortholog in *Ostreococcus tauri*. Nonetheless, some orthogroups containing genes in both species were found to be marked only in *Arabidopsis thaliana*. For example, the carotenoid oxygenase gene (CCD7 – *AT2G44990*) is H3K27me3 marked in *Arabidopsis thaliana*, while its *Ostreococcus tauri* orthologs *ostta01g04480*, *ostta03g03900* and *ostta03g05630* are not marked, Figure 12E.

Summing up, most H3K27me3 marked genes are species specific, reflecting substantial divergence in the regulatory targets of this epigenetic mark between *Arabidopsis thaliana* and *Ostreococcus tauri*. However, the striking conservation in the marking of transcription factors acting as key master regulators of gene expression indicates that while the specific downstream targets of H3K27me3 have diversified, its role in modulating higher-order regulatory nodes remains evolutionarily conserved.

## Discussion

Polycomb Group and Trithorax complexes play central roles in eukaryotic gene regulation both in animals^49^ and plants^50^, specifically in temperature response and acclimation. Nevertheless, their role as transcriptional key regulators in chlorophyte marine phytoplankton remained to be explored. Single copy genes were identified in the *Ostreococcus tauri* genome for most Polycomb Repressive Complex 2 and Trithorax components responsible for H3K27me3 and H3K4me3 deposition, respectively, highlighting its suitability as a model for chlorophyte marine phytoplankton due to its low genomic redundancy. Interestingly, the presence of two genes encoding orthologs for Suppressor of zeste 12 (Su[z]12) suggests the existence of different PRC2 variants in *Ostreococcus tauri*, which could be involved in distinct biological processes, similar to PRC2 variants in *Arabidopsis thaliana* containing VERNALIZATION2 (VRN2), EMBRYONIC FLOWER 2 (EMF2) or FERTILIZATION-INDEPENDENT SEED 2 (FIS2)^51^. No orthologs for the different components of the Polycomb Repressive Complex 1 responsible for H2Aub deposition were found in the *Ostreococcus tauri* genome, indicating H3K27me3 dynamics can be independent from H2Aub and that the coexistence of both marks could only be necessary in the green lineage for the response and acclimation to specific dynamics of environmental signals absent in the ocean. The lack of orthologs for the PRC1 components VAL1/2^32^ involved in recruiting PRC2 for the establishment of H3K27me3, indicates that other mechanisms involving, for example, transcription factors containing the EAR domain must be the ones used in *Ostreococcus tauri*^52^.

The size of *Ostreococcus tauri* cells, being the smallest known free-living eukaryote, suggests the presence of a massive compaction in its genome^22^. Nevertheless, H3K27me3 does not seem to play a primary role in overall genome compaction as it only occupies, under standard growth conditions, 15% of the entire genome length, a smaller proportion when compared to other organisms such as *Arabidopsis thaliana*. The potential role played by H3K27me3 in mediating local as well as long range genomic interactions in *Ostreococcus tauri* remains to be explored by integrating our results with Hi-C data^53^. H3K27me3 genome-wide distribution was only marginally associated to transposons resembling that of *Arabidopsis thaliana* occupying mainly the TSS and entire bodies of repressed genes distributed across the different chromosomes. H3K27me3 was found only significantly accumulated in the first half of chromosome 2 and the entire chromosome 19. These chromosomes have been associated with sexual reproduction and viral defense respectively^35,36^ .

H3K27me3 and H3K4me3 were found to be mutually exclusive, with H3K4me3 being associated with highly expressed genes. H3K4me3 was found occupying specifically the TSS of genes distributed across the entire genome, only being depleted from chromosome 19 and only slightly accumulating significantly in chromosome 20. Genes associated to specific biological processes such as ribosome biogenesis and assembly, translation initiation and photosynthesis were found to be H3K4me3 marked. Although a slight gradual, but significant, increase was observed in H3K4me3 levels at rising temperatures no significant effect was detected in the corresponding gene expression.

In contrast, the central role of H3K27me3 in regulating temperature acclimation in plants and animals was also found in *Ostreococcus tauri*, with mainly progressive (dose-dependent) increases at rising temperatures significantly associated with gene repression. This suggests quantitative non-binary H3K27me3 dynamics at the level of the entire cell culture that could result from a binary H3K27me3 marked / non-marked gene state at the level of individual cells. Although at the level of individual cells, H3K27me3 regulation over genes would be “digital” involving binary states OFF (H3K27me3 marked) / ON (H3K27me3 non-marked), the increasing proportion of individual cells with H3K27me3 marked genes at rising temperatures would result, after averaging, in an apparent gradual quantitative “analog” H3K27me3 regulation mode at the level of the entire cell culture. This integration between analog and digital modes of gene regulation exerted by H3K27me3 is consistent with the one described for *Arabidopsis thaliana*^54^.

H3K27me3 marked genes were significantly enriched in specific biological processes indicating a regulatory role of this mark. Genes encoding motor proteins such as dyneins, kinesins and myosins were found H3K27me3 marked at high temperatures. Whereas H3K27me3 levels did not change significantly with temperature in genes codifying for dyneins, at lowering temperatures a gradual removal of H3K27me3 was detected in genes codifying for kinesins and myosins. This was concomitant with a gradual increase in expression in the corresponding genes, Figure 13. Similarly, genes involved in microtubule and cortical actin network formation exhibited decreasing H3K27me3 levels and increasing expression at lowering temperatures. This suggests that, at low temperatures, cellular architecture is remodeled to activate a dual-range vesicle trafficking: long-range transport mediated by kinesins along microtubules, combined with short-range vesicle delivery via cortical acting networks by myosins, potentially leading to secretion, Figure 13. Lipid biosynthesis was also found to be partially regulated by H3K27me3, suggesting that these vesicles may contain lipids among other compounds. Genes involved in polyketide biosynthesis, lipid transport and biosynthesis, such as the Enoyl-ACP Reductase, were detected as H3K27me3 marked at high temperatures, whereas at lowering temperatures this mark was removed and gene expression activated, Figure 13. The activation of lipid vesicle trafficking at low temperature was validated using microscopy imaging and lipid staining. Lipid vesicle secretion may seem a counterintuitive strategy under environmental stress such as a low temperature. Nonetheless, this process has been suggested to be beneficial in defending cells against viruses since vesicle containing external membrane receptors recognized by viruses may act as baits capturing viruses and preventing them from infecting and killing cells^55^.

**Figure 13.**
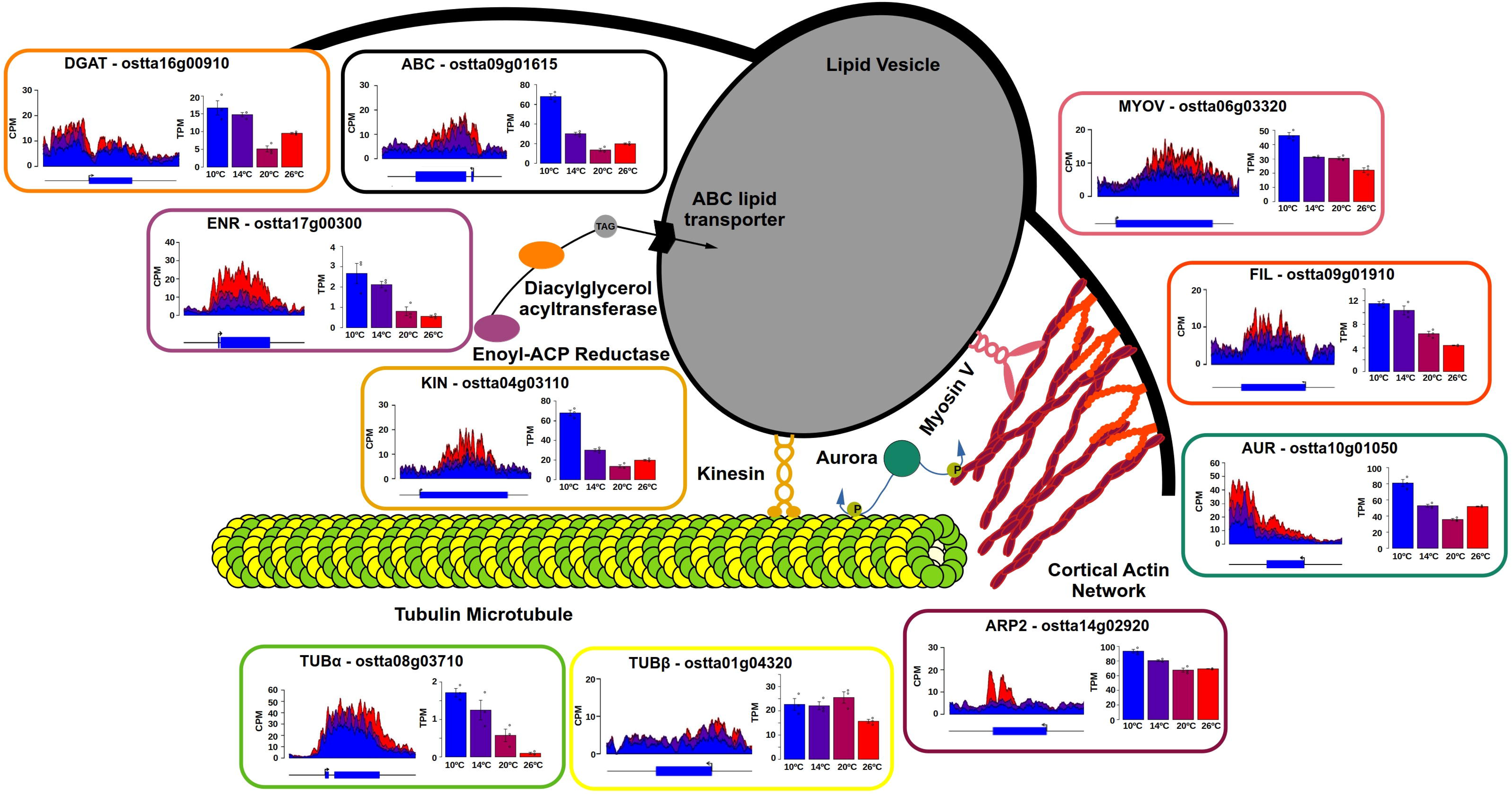
Model of the biological processes regulated by H3K27me3 in temperature acclimation. Graphical representation of the expression level increase and H3K27me3 ChIP signal decrease at lowering temperature of genes involved in microtubule and cortical actin network formation and activation, tubulin alpha (TUBα) and beta (TUBβ), Actin Related Protein 2 (ARP2), MORN-Filamin-Invasin_D3 domain-containing protein (FIL) and Aurora (AUR); long-range microtubule based motor proteins, kinesin (KIN) and short-range cortical actin network delivery motor proteins myosin V (MYOV); and lipid biosynthesis and transport, Enoyl-ACP Reductase (ENR), Diacylglycerol Acyltransferase (DGAT) and ABC transporter (ABC). For each gene, H3K27me3 ChIP signal is represented on the left measured using CPM (Counts Per Million) and gene expression is presented on the right measured using TPM (Transcripts Per Million). Blue represents 10°C, purple 14°C, magenta 20°C and red 26°C.

H3K27me3 is also expected to take part in the regulation of the yet uncharacterized sexual cycle in *Ostreococcus tauri*^36^. Specifically, H3K27me3 was found marking genes involved in the pachytene stage of meiosis regulating synaptonemal complex formation to ensure proper chromosome pairing and recombination. Although reduced H3K27me3 levels were found at low temperatures at these genes, no significant expression activation was detected. This suggest that low temperature only primes these genes for activation and other signals would be necessary for fully gene expression activation, possibly light as reported for other microalgae, to induce the sexual reproductive cycle in *Ostreococcus tauri* ^56,57^.

In order to obtain insights into the evolution and diversity of the regulatory roles of H3K27me3 in photosynthetic eukaryotes, its gene targets in *Ostreococcus tauri* were compared, using orthology relationships, with those previously described for *Arabidopsis thaliana*. Little conservation between both species was found among H3K27me3 targets, except for the transcription factor families MADS-box, WRKY and AP2, with orthologous genes marked in both species. In *Arabidopsis thaliana*, members of these families have been shown to play central roles in regulating key processes in response to temperature associated with sexual reproduction, such as the flowering transition and floral development^58^, as well as immune system response^59^. Our results show that H3K27me3 regulates meiosis and lipid vesicle trafficking which could be considered to be part of the sexual and immune system response in *Ostreococcus tauri*. This suggests that this epigenetic mark regulates these fundamental biological processes, sexual reproduction and immune system, in these two distant species in the green lineage through the same key transcription factor families, although the specific intermediary mechanisms differ. This indicates that, although the specific downstream targets of this epigenetic mark have diversified during evolution, both the biological processes and their higher order regulatory nodes remain evolutionarily conserved.

## Methods

### Culture conditions and sample collection

Cultures of the *Ostreococcus tauri* sequenced strain RCC4221 were grown in flasks on orbital shakers within phytotrons under constant light (100 μE m^-2^s^-1^) at 10°C, 14°C, 20°C and 26°C, with three independent biological replicates per condition. The growth medium consisted of sterilized artificial sea water (ASW)^60^ supplemented with nitrates, phosphates, trace metals and vitamins as described in ^3^. Samples were collected one week after the initial inoculation at a starting concentration equivalent to 6 mg L^-1^ chlorophyll.

### Chromatin fixation, cell disruption, fragmentation, immunoprecipitation, DNA purification and sequencing

Approximately 5×10^9^ cells were harvested by centrifugation for 4 min at 7000 x g and 4°C. The supernatant was discarded and the cells were washed with phosphate-buffered saline (PBS) 1X solution.

Chromatin fixation was performed by resuspension in Cross-linking solution (1% Formaldehyde dissolved in PBS 1X) and incubation in a vacuum desiccator for 10 min at room temperature (RT). Chromatin cross-linking was ceased by adding glycine to a final concentration of 0.125 M and incubating for 10 min at RT followed by three cold PBS 1X washes.

Cells were disrupted after resuspension in Extraction Buffer 1 (0.4 M sucrose, 10 mM Tris-HCl pH 8, 10 mM MgCl_2_, 5 mM β-mercaptoethanol, 0.1 mM PMSF inhibitor) with a single freeze-thaw cycle using a mortar and liquid nitrogen. Next, samples were centrifuged for 10 min at 11000 x g at 4 °C and resuspended in Extraction Buffer 2 (0.25 M sucrose, 10 mM Tris-HCl pH 8, 10 mM MgCl_2_, 1% Triton X-100, 5 mM β-mercaptoethanol, 0.1 mM PMSF inhibitor, 1X Protease Inhibitor Cocktail) and incubated on ice for 10 min and centrifuged again under the same conditions to remove the supernant. This last step was repeated once again and the samples were resuspended in Low Salt Buffer (150 mM NaCl, 0.1% SDS, 1% Triton X-100, 2 mM EDTA, 20 mM Tris-HCl pH 8) supplemented with 1X Protease Inhibitor Cocktail. Samples were finally divided into three different aliquots and stored at –80 °C.

Fixed chromatin fragmentation was carried out by sonication in a Bioruptor Pico device (Diagenode) using 15 cycles of 30 s ON / 30 s OFF. Subsequently, the samples were incubated on a wheel at 4°C for 30 min, followed by centrifugation at 11000 x g for 10 min at 4 °C. Next, the supernatant was collected for further processing. Sonication efficiency in generating fragments around 200-1000bp was tested using sonicated chromatin, reverse cross-linking and gel electrophoresis to determine DNA size.

Chromatin immunoprecipitation was performed by adding 2 μl of antibody (anti-H3K27me3 or anti-H3K4me3 Diagenode) to 300 μl of each fragmented sample and incubating them overnight on a wheel at 4 °C. 30 μl of sonicated chromatin were used as INPUT samples and stored at –20 °C until required. Protein A sepharose beads (30 μl/sample) were hydrated in Low Salt buffer by rotation on a wheel for 30 min at 4 °C, washed 3 times with the same buffer and centrifuged for 1 min at 100 x g. Next, the beads were blocked by resuspension in a blocking buffer (950 μl Low Salt buffer and 50 μL of sheared salmon sperm DNA 10 mg/ml) and incubated overnight on a wheel at 4 °C. The following day, the beads were washed 3 times with Low Salt buffer and then mixed with their corresponding samples. Immunocomplexes were settled under a 2 hour and a half incubation on a wheel at 4 °C. The immunocomplexes were washed four times using the following buffers sequentially: Low Salt, High Salt (500 mM NaCl, 0.1% SDS, 1% Triton X-100, 2 mM EDTA, 20 mM Tris-HCl (pH 8)), LiCl (0.25 M LiCl, 1% NP-40, 1% sodium deox-ycholate, 1 mM EDTA, 10 mM Tris-HCl (pH 8)) and TE (10 mM Tris-HCl (pH 8), 1 mM EDTA). Each wash was performed followed by a 5 min incubation on ice and a centrifugation of 1 min at 100 x g. Immunocomplexes elution from the beads was achieved by adding 250 μl of freshly prepared Elution buffer (1% SDS, 0.1 M NaHCO3) to the samples and incubating them for 30 min at 65 °C with gentle agitation. After centrifugation for 5 min at 5000 x g, the supernatants were passed to new eppendorfs. These immunoprecipitated (IP) samples were mixed with 10 μl NaCl 5 M. INPUT samples were defreezed and mixed with 220 μl Elution buffer and 10 μl NaCl 5 M and chromatin cross-linking was reversed by overnight incubation at 65 °C.

Sequencing libraries were generated following manufacturer’s instructions and sequencing was performed on an Illumina NextSeq500 sequencer for 3 replicates for each temperature conditions including IP and INPUT samples. Approximately 9 million 75 nt long single end reads were generated for each sample.

### ChIP-seq data analysis

ChIP-seq data were analyzed using our pipeline MicroAlgae RNA-seq and Chip-seq AnalysiS (MARACAS)^61^. Specifically, quality control of the sequencing data was performed using the software package FASTQC. The *Ostreococcus tauri* genome sequence and annotation v3.0 (https://phycocosm.jgi.doe.gov/Ostta4221_3/Ostta4221_3.home.html) were used as reference genome. Reads were mapped to the reference genome with bowtie2^62^. SAM (Sequence Alignment Maps) and BAM (Binary Alignment Maps) files were processed using the software tool SAMtools^63^. Significant ChIP signal peaks or ChIP enriched regions were identified for each temperature replicate comparing the corresponding ChIP and INPUT samples using the software package MACS2 (Model-based Analysis of ChIP-seq)^64^. These ChIP signal peaks were stored in BED (Browser Extensible Document) format that were processed using the software toolboxes BEDTools^65^ and deepTools^66^. For each temperature condition ChIP signal peaks identified consistently in all three replicates were determined using intersectBed. The complete set of consensus ChIP signal peaks identified in at least one temperature condition were computed using mergeBed. Normalized ChIP signal levels were estimated over the set of consensus peaks as CPM (Counts Per Million of mapped Reads) by first computing raw read counts using intersectBed and dividing by the millions mapped reads in each sample.

Genome wide distribution of ChIP signal peaks and gene targets were determined using the Bioconductor R packages ChIPpeakAnno^67^ and the genomic R package developed by our group for *Ostreococcus tauri* TxDb.Otauri.JGI available in our GitHub repository (https://github.com/fran-romero-campero/AlgaeFUN/tree/master/packages/txdb_packages). Principal Components Analysis (PCA) and Hierarchical Clustering (HC) were performed using the R package FactoMineR^68^. Peaks with significant changes in ChIP signal between temperature conditions or Differentially Occupied Regions (DORs) were identified according to a moderated t test implemented in the R package limma^69^ according to an adjusted p-value threshold of 0.01 and a fold change cutoff of 1.5. The R code for this analysis is available from the GitHub repository ECTOR (https://github.com/fran-romero-campero/ECTOR).

### RNA extraction, purification and sequencing

For each temperature replicate, 50 mL of culture were collected for RNA extraction. Cells were washed with PBS using centrifugation for 1 min at 13,000 × g and 4°C. After supernatant removal, cells were immediately flash frozen in liquid nitrogen and stored at −80 °C. RNA extraction and purification were performed as described in ^3^. Sequencing libraries were generated following manufacturer’s instructions and sequencing was performed on an Illumina NextSeq500 sequencer. Approximately 15 million 100 nt long single end reads were generated for each one of the three replicates for each temperature condition.

### RNA-seq data analysis

MicroAlgae RNA-seq and Chip-seq AnalysiS (MARACAS)^61^ was used to analyze our RNA-seq data. Briefly, FASTQC was used to check the high quality of the sequencing data, reads were mapped to the reference genome with HISAT2^70^, transcript assembly and gene expression estimation measured as transcripts per million of mapped reads (TPM) and raw counts were performed using StringTie2^71^. PCA and HC were performed using the R package FactoMineR^68^. Differentially expressed genes were determined using the negative binomial test implemented in the Bioconductor R package DESeq2^72^ according to an adjusted p-value threshold of 0.05 and a fold change cutoff of 2. The R code for this analysis is available from the GitHub repository ECTOR (https://github.com/fran-romero-campero/ECTOR).

### Functional enrichment analysis, transposable elements identification and evolutionary analysis

Gene ontology enrichment analysis over the different gene sets of interest were performed using the web app AlgaeFUN^61^ (https://greennetwork.us.es/AlgaeFUN/) based on the Bioconductor R package clusterProfiler^73^ and the annotation package for *Ostreococcus tauri* org.Otauriv5.eg.db developed by our group based on recent annotation updates^26^ available from the GitHub repository https://github.com/fran-romero-campero/OSTIGO/blob/main/org.Otauriv5.eg.db.zip.

Transposable elements were identified using the software tools RepeatMasker and RepeatModeler2^74^ with default parameters.

Orthogroups for *Arabidopsis thaliana* and *Ostreococcus tauri* evolutionary comparative analysis were obtained from the web app PharaohFUN^75^ (https://greennetwork.us.es/PharaohFUN/) which is based on the software platform OrthoFinder^76^.

### Transmission electron microscopy

For each sample, 100 mL of fresh *Ostreococcus* cultures at a concentration equivalent approximately to 10 mg L^-1^ chlorophyll were centrifuged for 5 min at 10000 x g and RT. The resulting pellet was washed once with fresh medium (ASW supplemented as described above), resuspended in Fixing solution (Glutaraldehyde 2% and Paraformaldehyde 4% diluted in fresh medium) and incubated on ice for 1 h. Next, the pellet was washed three times with fresh medium, resuspended in 1% OsO_4_ and incubated for 1 h at 4°C. After three washing cycles in fresh medium (20 min each one), samples were immersed in 2% uranyl acetate, dehydrated through a gradient acetone series (50%, 70%, 90% and

100%) and embedded in Spurr resin^77^. Blocs were obtained by polymerization at 70°C for 8 h. Thin sections were cut with a diamond knife in an ultramicrotome (Leica UC7) and examined with a transmission electron microscope (Zeiss Libra 120) operating at 80 kV. The electron microscopy images were taken at different magnifications with an EMCCD camera (TRS 2k x 2k).

### Nile Red staining and confocal microscopy

For each sample, fresh Ostreococcus culture comprising approximately 10^8^ cells was mixed with 50ul of Nile Red dye (500 ug/ml) to a final volume of 300 ul. The mixture was incubated for 30 minutes at 37°C in darkness, followed by centrifugation for 5 min at 10000 xg at RT. This process was repeated one more time and the supernatant was discarded. DMSO was used as negative control.

Confocal microscopy images of *Ostreococcus* cells stained with Nile Red were acquired using a spectral Laser Scanning Confocal Microscope (Olympus FLOUVIEW FV3000). Excitation was performed with a 488 nm laser. Emission signals were detected within the green channel (530 to 575 nm, gain 480V) for Nile Red and red channel (650 to 750 nm, gain 400V) for chloroplasts.

## Data and Code availability

ChIP-seq and RNA-seq data generated in this study are freely available from the Gene Expression Omnibus database in the superseries identified with the accession number GSE237488 (https://www.ncbi.nlm.nih.gov/geo/query/acc.cgi?acc=GSE237488).

The complete R code for all the analysis perfomred in this study is available from the GitHub repository ECTOR (https://github.com/fran-romero-campero/ECTOR).

## Acknowledgements

We would like to acknowledge Myriam Calonje for guiding Christina Arvanitidou in the development of the protocol used in this study for generating ChIP-seq data and for critical reading of this manuscript. We would like to thank Eloisa Andújar and Mónica Pérez from the CABIMER Genomics Unit for their assistance with high-throughput sequencing, Alicia Orea from the IBVF Microscopy Service for her help with confocal microscopy imaging and Juan Luis Ribas-Salgueiro for his cooperation in the generation of the samples and images for transmission electron microscopy.

## Author contributions

C.A. and M.E.G.G. performed wet lab experiments. C.A. generated all ChIP-seq and RNA-seq data. C.A. and F.J.R.C. performed multiomics data analysis and integration. C.A., M.E.G.G. and M.G.-G. carried out samples processing for transmission electron microscopy, nile red staining and confocal microscopy. M.R.G., F.J.R.C. and C.A. carried out the evolutionary comparative analysis. F.J.R.C., C.A., M.G.G. and F.C. designed all experiments, interpreted the results and wrote the manuscript. All authors read and approved the final manuscript.

---

**Supplementary Figure 1.**
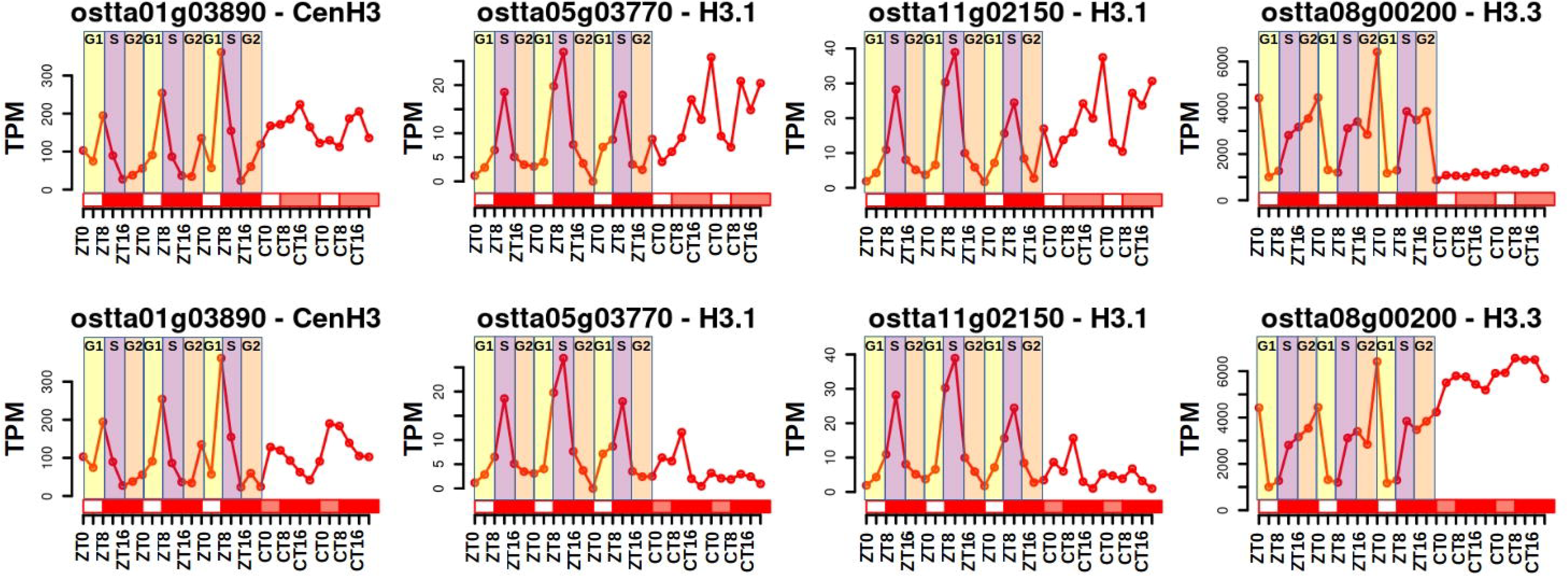
Supports Figure 1. Expression profiles for genes encoding Histone 3 CenH3 centromeric varian, H3.1 canonical variant and H3.3 variant in *Ostreococcus tauri* measured as TPM (Transcripts per Million) under short day conditions and constant light, top, and constant darkness, bottom. Gl cell cycle phase is marked in yellow, S cell cycle phase is marked in purple and G2/M cell cycle phase is marked in orange. ZTN, Zeitgeber time N, marks the time point N hours after dawn (lights on). CTN, circadian time N, denotes the time point N hours after subjective dawn.

